# Biasing Conformational Sampling in AlphaFold 3 and Boltz-2 via Pair Representation Scaling

**DOI:** 10.64898/2026.01.23.701250

**Authors:** Shosuke Suzuki, Toshiyuki Amagasa

**Affiliations:** Graduate School of Science and Technology University of Tsukuba, Tsukuba, 305-8573, Japan; Center for Computational Sciences University of Tsukuba, Tsukuba, 305-8577, Japan

**Keywords:** protein conformational sampling · structure prediction · deep learning

## Abstract

Deep learning has transformed protein structure prediction, yet most systems return a single dominant conformation with little control over alternative functional states. We introduce pair representation scaling, an inference-time method that biases conformational sampling in diffusion-based structure predictors by multiplying the latent pair representation by a single scalar before the Pairformer trunk, without retraining, an auxiliary model, or a second forward pass. On 86 two-state targets spanning domain motions and membrane transporters, scaling broadens the conformational ensembles of both AlphaFold 3 and Boltz-2 and recovers alternative states that default inference misses, most strongly in AlphaFold 3, where the gains extend even to targets deposited after the training cutoff. It approaches the alternative-state recovery of alignment-based sampling methods, and the benefit persists even without a multiple sequence alignment. The predicted distance distributions show that scaling shifts the encoded two-state distribution toward the experimentally observed alternative state, a directed modulation rather than arbitrary perturbation. Pair representation scaling is an interpretable, low-cost handle on the conformational ensembles of deep learning structure predictors.

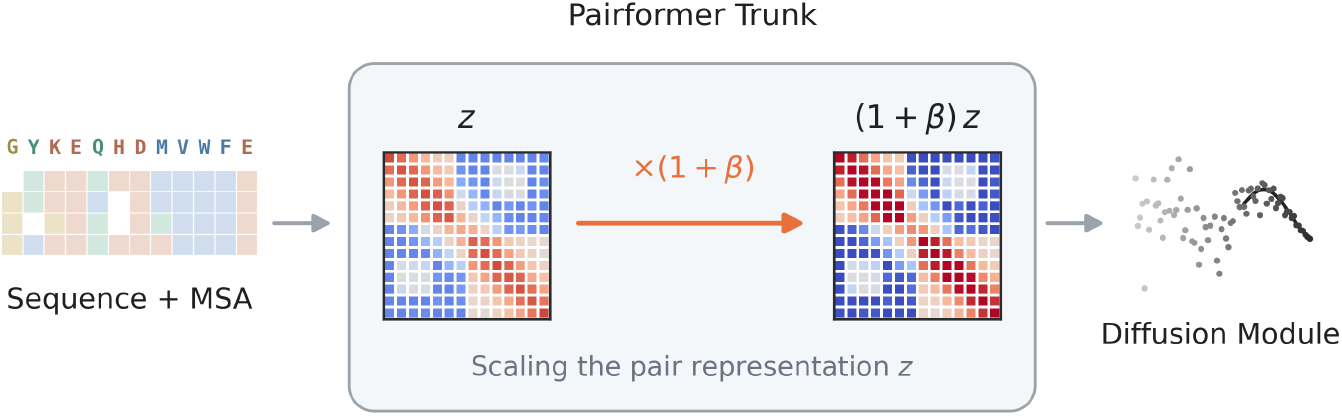

**Pair representation scaling:** A query sequence and its multiple sequence alignment enter the Pairformer trunk, where the latent pair representation *z* is rescaled by a single global scalar to (1 + *β*) *z* and the diffusion module then generates a structure. The alignment and the trained weights are unchanged, and the same operation applies in AlphaFold 3 and Boltz-2. The scalar *β* is a single explicit handle: sweeping it broadens the sampled ensemble toward alternative states, and its effect can be read out directly inside the network.

## 1 Introduction

Deep learning has transformed the prediction of protein three-dimensional structures from amino acid sequences, in particular through AlphaFold 2 [1]. While such models achieve near experimental accuracy, they typically return a single dominant conformation for each sequence. In reality, many proteins are dynamic systems that interconvert between distinct structural states to perform their biological functions [2]. Understanding how deep learning models represent this conformational diversity, and how alternative states can be accessed in a steerable way, remains an open problem in structural biology and biophysics [3, 4]. Conformational heterogeneity also shapes molecular recognition and allostery, motivating long-standing efforts to represent proteins as ensembles rather than single structures. Traditional approaches rely on molecular dynamics or enhanced sampling to generate ensembles, which can be computationally demanding and require system-specific expertise to parameterize.

Recent work has shown that modifying the multiple sequence alignment (MSA) can reveal alternative structural states [5, 6, 7, 8, 9, 10]. Approaches that cluster or selectively subsample sequences, such as AF-Cluster [11], aim to isolate weaker coevolutionary signals associated with distinct conformational preferences. These results suggest that MSAs encode both dominant structural information and subtler evolutionary signals for alternative conformations. In practice, however, such MSA-based procedures act entirely through the input alignment, so they cannot run without one and their cost grows with its depth. In parallel, inference-time strategies act on the network’s internal computation rather than on its input. Dropout-based sampling reactivates the trained dropout layers during inference to inject stochastic noise [12], and entropy-guided folding modifies the intermediate representations of AlphaFold 2 by back-propagating a distogram entropy objective [13]. Complementary studies have begun to exploit AlphaFold-derived ensembles directly for thermodynamic and functional predictions, for example to model enzyme thermostabilization or kinase activation-loop conformations relevant for inhibitor binding [14, 15], and a broader literature situates these efforts within the open question of how AlphaFold relates to protein dynamics and allostery [16]. However, these approaches reshape the input alignment, inject generic stochasticity, or optimize an auxiliary objective over the representations, and none offers a single, interpretable parameter for shifting the relative weights of alternative conformations.

Newer architectures such as AlphaFold 3 [17] and Boltz [18, 19] streamline MSA processing and organize the network trunk around a Pairformer module that feeds a diffusion-based coordinate generator [20]. In these models, sequence features are integrated into a latent pair representation that aggregates coevolutionary and geometric information and serves as the main input to the structure module. This separation between sequence, pairwise, and coordinate representations creates a natural interface for perturbations of the network’s internal representations. In particular, the pair representation can be viewed as an effective residue–residue coupling field that mediates how evolutionary constraints are translated into structural geometry.

Here we introduce pair representation scaling, an inference-time method that rescales the latent pair representation by a single scalar *β* to modulate conformational sampling. The operation is a single multiplication of one internal representation, applied without retraining and without altering the input alignment, and because that representation is shared across recent diffusion-based predictors the same operation applies to AlphaFold 3 and Boltz-2 without modification. Because it acts on an internal representation rather than the alignment, the rescaled signal can be read out directly, the operation stays usable when the alignment is removed, and it composes with alignment-based sampling. Benchmarked across 86 two-state targets against alignment-based and inference-time baselines and the ensemble generator BioEmu, scaling expands the sampled conformational ensemble in both predictors, and the predicted distance distributions show that the broadening is directed toward the experimentally observed alternative state.

## 2 Results

### 2.1 Pair representation scaling

AlphaFold 3 and Boltz-2 share a trunk in which a Pairformer refines single and pair latent representations that condition a diffusion-based structure generator [17, 19]. The pair representation is the principal route through which evolutionary and structural constraints reach the structure module, and because it is shared across recent diffusion-based predictors, it offers a single interface for an intervention that applies to either architecture without modification.

We define pair representation scaling, an inference-time operation that multiplies the latent pair representation by a single scalar at the input to the Pairformer. Let *z_ij_* ∈ R^d^*^P^* denote the pair embedding for residues (*i, j*) produced by the conditioning trunk. We apply

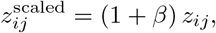

with the scaling applied at each recycling iteration. The input sequence, the MSA, and the trained weights all remain unchanged. The factor (1 + *β*) adjusts the effective magnitude of the pair representation relative to the value the network would otherwise use. Because pair features enter the Pairformer through additive and projection pathways in its attention and transition blocks, rescaling *z_ij_* changes the relative contribution of pair-derived signal to the trunk and, in turn, to the conditioning passed to the diffusion module.

The operation is a single element-wise multiplication of a tensor already produced during standard inference, so it adds negligible computation on top of a forward pass, and applying it to a predictor is a small change to the inference code. We apply the same operation to AlphaFold 3 and to Boltz-2, sweeping *β* over a fixed symmetric range that perturbs the generated ensemble while preserving stable inference (Section 5.1.1).

### 2.2 Benchmark across conformational-change types and the training cutoff

We assembled a benchmark of 86 two-state targets, namely 39 domain-motion targets and 47 membrane transporters that interconvert between inward-facing and outward-facing states (Section 5.2). For each predictor we report the benchmark in three groups, the domain motions and the transporters whose reference structures predate that predictor’s training cutoff, and an after-cutoff group of targets whose structures were released after it. The after-cutoff group isolates whether a gain depends on the alternative state having been seen during training, a question raised in particular for the alternative conformational states of solute carrier membrane proteins, where memorization bias has been documented to favour one state over the other [21], one of several blind spots that alternative folds expose in these predictors [22]. Because the predictors have different cutoffs, this group holds 23 targets for AlphaFold 3 and 13 for Boltz-2 (Section 5.2.3). We compared pair representation scaling with default inference, with three established perturbations of the input alignment, namely subsampling [6], random masking of MSA columns [10], and clustering [11], with inference-time dropout [12], with the MD-conditioned mode of Boltz-2 [19], and with the structure-ensemble generator BioEmu [23]. The three predictors differ in how directly their training targets conformational ensembles. AlphaFold 3 is trained on static PDB structures [17], Boltz-2 additionally on molecular-dynamics ensembles [19] from MISATO [24], ATLAS [25], and mdCATH [26], and BioEmu in three stages specifically to emulate equilibrium ensembles, combining pretraining on the AlphaFold database, molecular-dynamics trajectories reweighted with Markov state models, and fine-tuning on the MEGAscale stability dataset [23]. We include BioEmu as a reference generator rather than a target-matched baseline, because each predictor’s groups follow its own training cutoff over a partly different set of targets. It shows what direct ensemble training recovers on the same benchmark. We applied the method to both AlphaFold 3 and Boltz-2, alone and combined with the alignment-based perturbations, and report three metrics (Fig. 1). The per-state success rate is the fraction of reference states for which the ensemble contains a model within 2 Å. The worst-case minimum RMSD is the larger of the two minimum distances to the reference states, and measures recovery of the harder state. The fill ratio measures how completely the ensemble covers the path between the two states.

**Figure 1.**
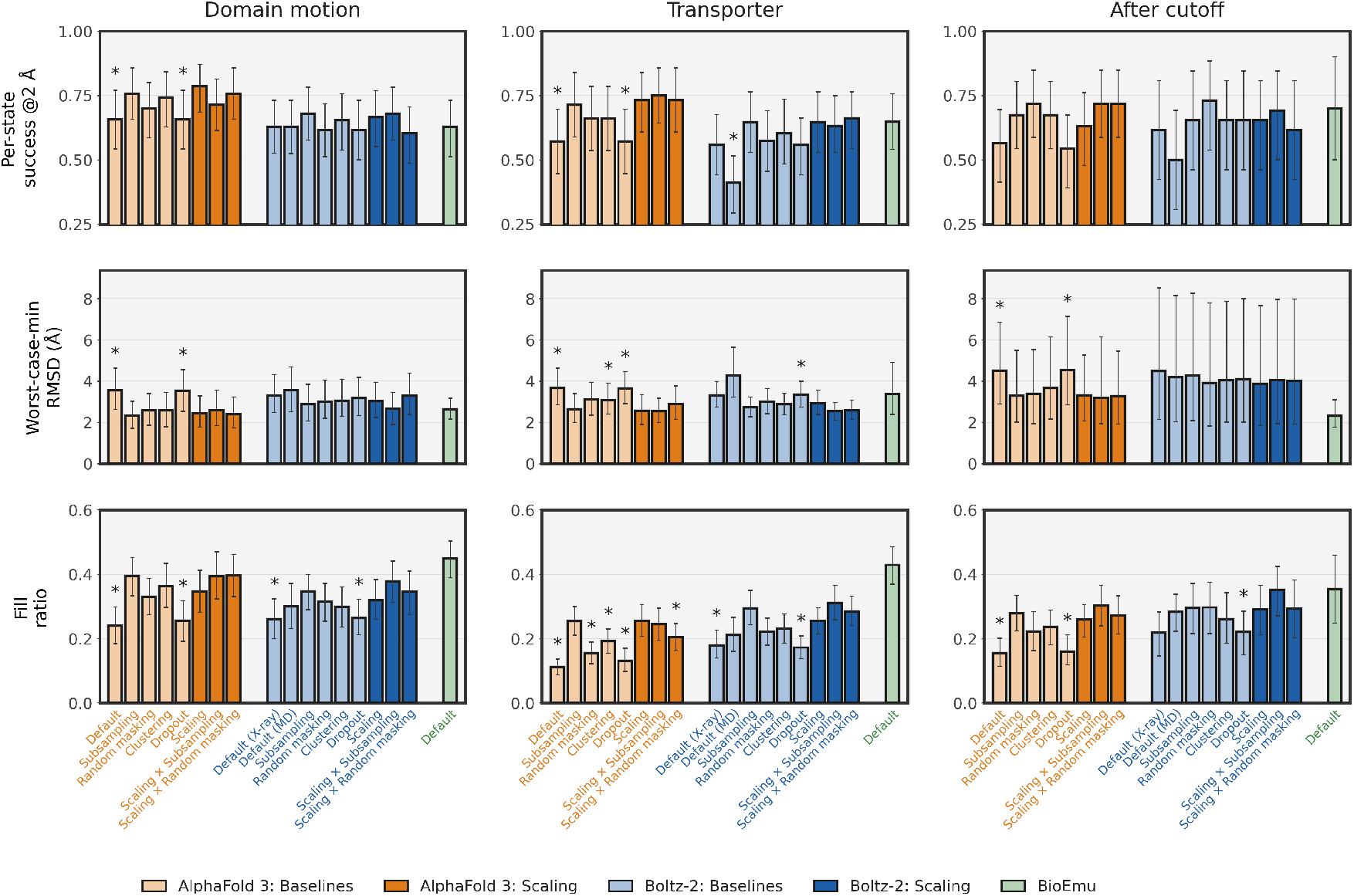
Benchmark across conformational-change types and the training cutoff. Per-state success rate, worst-case minimum RMSD, and fill ratio for the domain-motion, transporter, and after-cutoff groups. Per-state success rate and fill ratio are higher-is-better and worst-case minimum RMSD is lower-is-better, and within each row the three category panels share one vertical scale. Group sizes for the domain-motion, transporter, and after-cutoff groups are AlphaFold 3 (35, 28, 23), Boltz-2 (39, 34, 13), and BioEmu (39, 37, 10). Bars are grouped by predictor. Scaling and its combinations with MSA subsampling and random masking are in the saturated colour and the baselines in the lighter tint. Bars are target means and error bars are 95% bootstrap confidence intervals. An asterisk on a bar marks a method that pair representation scaling significantly outperforms (paired Wilcoxon signed-rank test, Holm-corrected within each predictor, *p <* 0.05).

In AlphaFold 3, pair representation scaling improved all three metrics across all three groups, with significant reductions in worst-case minimum RMSD over default inference in every group. The per-state success rate rose from 0.60 under default inference to 0.73 under scaling across the 86 targets, with significant gains in both the domain-motion and transporter categories against default inference after Holm correction (paired Wilcoxon signed-rank test). The fill ratio rose in both categories and more than doubled on transporters. Among the baselines, the alignment-based perturbations were the closest competitors, which weaken the dominant alignment signal, and scaling matched the strongest of them.

In Boltz-2, the same operation broadened the conformational ensemble with a smaller but consistent effect. Scaling improved all three metrics over default inference in all three groups, raising the per-state success rate by up to 0.09 on transporters and lowering the worst-case minimum RMSD by 0.28 to 0.60 Å. The fill-ratio gain reached significance in all three groups after Holm correction (*p <* 0.001 in the two pre-cutoff groups and *p* = 0.04 after the cutoff), and on transporters scaling improved significantly over inference-time dropout in worst-case minimum RMSD and over the MD-conditioned mode in per-state success. Against default inference the per-state and worst-case gains were consistent in direction but did not cross the significance threshold (*p* ≈ 0.06 for the worst-case improvement in the two pre-cutoff groups). The improvement that reached significance against default inference was therefore the fill ratio, with per-state success and worst-case recovery shifting in the same direction.

Combining scaling with MSA subsampling helped most in Boltz-2. The combination lowered the worst-case minimum RMSD below default inference in both categories, which scaling alone did not reach (*p <* 0.001), and raised the fill ratio above scaling alone (*p <* 0.01). Its per-state success rate rose over default inference on transporters (*p* = 0.02) but not on domain motions. In AlphaFold 3, where scaling alone was already strong, the combination did not improve on it overall, adding to the fill ratio on domain motions but leaving the worst-case minimum RMSD comparable and the per-state success rate mixed, slightly higher on transporters and slightly lower on domain motions. The combination therefore recovers the harder Boltz-2 metrics that the internal perturbation alone leaves unchanged, whereas in AlphaFold 3 scaling alone suffices.

BioEmu, trained directly on equilibrium ensembles, produces the broadest fill ratio of the three predictors (0.43 against 0.29 for AlphaFold 3 under scaling). On the after-cutoff group the AlphaFold 3 worst-case minimum RMSD improved significantly, by 1.18 Å (95% bootstrap CI 0.47–2.03 Å) over its 23 targets, so the gains extend to targets whose alternative state was not available during training.

Measured as the fraction of the ensemble within 2 Å of each reference rather than as an extremum, the occupancy of the harder state rose under scaling in all three AlphaFold 3 groups and in two of the three Boltz-2 groups, while the occupancy of the dominant state fell in both predictors (Section S2).

### 2.3 Per-target recovery of the alternative state

The aggregate gain in the benchmark is the sum of per-target shifts in which a scaled ensemble reaches a reference state that default inference leaves unrecovered. Figure 2 shows this pattern for five representative targets in each predictor, three drawn from its training set and two with reference structures released after its training cutoff, spanning both soluble domain motions and membrane transporters. The per-target scatters for all 86 benchmark targets are provided in Section S5. For each target the default ensemble clusters close to one of the two reference states while leaving the other far from recovery, and scaling spreads the ensemble toward the second state. The predicted structures confirm the recovery, as the ensemble models nearest the two references reproduce both the dominant and the alternative conformation. The pattern holds across soluble domain motions, such as adenylate kinase and the glutamine-binding protein, and membrane transporters, such as the lactose permease and the nucleoside transporter, in both predictors. In every case the scaled ensemble overlaps the default cluster and widens it toward the alternative state, and the value of *β* that surfaces the alternative state differs in sign between targets.

**Figure 2.**
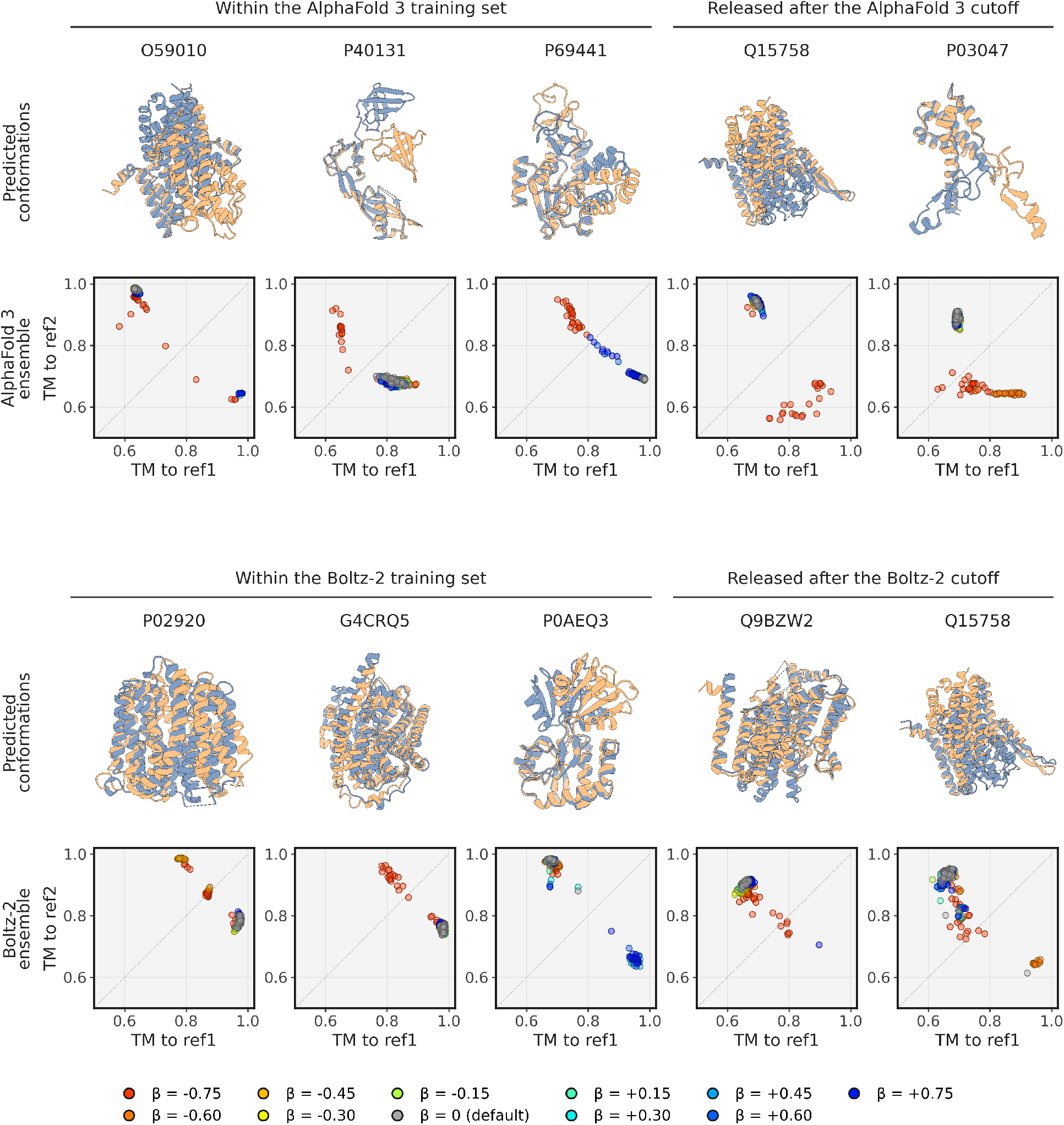
Per-target recovery of the alternative state. Five representative targets for AlphaFold 3 (upper half) and five for Boltz-2 (lower half), three from each predictor’s training set and two with reference structures released after its training cutoff. For every target the superposed structures are the two ensemble models closest to the two experimental reference states, coloured slate blue and tan, with transporters viewed along the membrane normal and soluble proteins along their principal axes. The scatter below each plots the TM-score to one reference against the TM-score to the other for all models, grey for default inference and coloured by *β* for scaling.

### 2.4 Structurally intermediate conformations

The benchmark and the per-target results describe a broadening of the ensemble toward the alternative reference state. As a further test of that direction, we asked whether the scaled ensemble also approaches other deposited conformations that lie structurally between the two reference states [9]. For each of the 39 domain-motion targets we retrieved every non-reference PDB entry cross-referenced from the target’s UniProt record and computed its TM-score to each reference. Entries with both TM-scores between 0.70 and 0.95 were taken as structurally intermediate, lying between the two end-states in structure space (Section 5.6). The criterion yielded 75 such structures across 13 of the 39 targets, with the remaining 26 contributing none.

Across the 75 structures, the best TM-score from the prediction ensemble was higher under scaling than under default inference in both predictors (per-intermediate paired Wilcoxon signed-rank test, *p* = 3 × 10*^−^*^5^ for AlphaFold 3 and *p* = 7 × 10*^−^*^4^ for Boltz-2). Because the 75 intermediates come from only 13 targets, we also aggregated the improvement per target and tested across the 13 (Fig. 3B). The gain held at the target level for AlphaFold 3 (11 of 13 targets improved, *p* = 0.01) but not for Boltz-2 (9 of 13, *p* = 0.25), so the Boltz-2 effect is directional rather than significant once the targets are the unit. The improvement is small in absolute TM-score (mean 0.014 in AlphaFold 3 and 0.007 in Boltz-2). An intermediate within a TM-score of 0.70 to 0.95 of both references is already close to any model that reaches either reference, so the headroom is limited. The smaller Boltz-2 gain reflects a higher starting point: its default inference already matches the intermediates more often than AlphaFold 3 (median best TM-score 0.93 against 0.91), leaving fewer to recover, consistent with its molecular-dynamics training giving broader default coverage. At the more stringent threshold of best TM *>* 0.90, AlphaFold 3 scaling reached nine more of these structures than default inference (McNemar exact test *p* = 0.004), and the threshold did not separate the two Boltz-2 conditions. Three illustrative targets show the same pattern at the per-target level (Fig. 3A).

**Figure 3.**
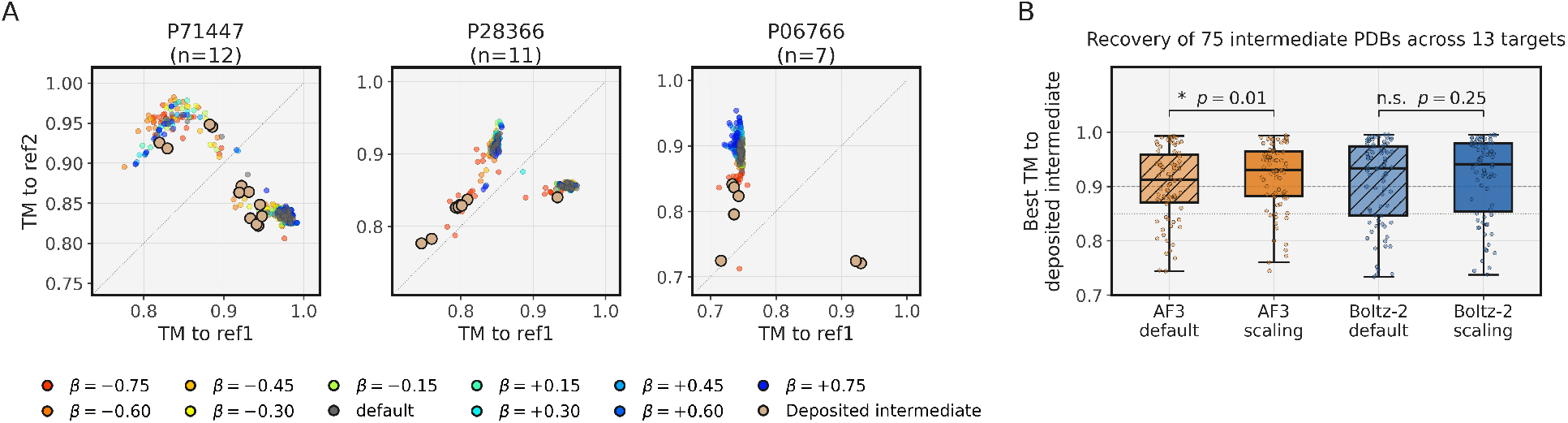
Structurally intermediate conformations. **(A)** The three domain-motion targets with the most deposited intermediate structures (P71447, P28366, and P06766). Each deposited intermediate is a filled disc, and the scaled AlphaFold 3 samples are coloured by *β* over a grey default-inference subsample. **(B)** Best ensemble TM-score to each of the 75 intermediate structures, as box-and-strip plots for the four conditions. Brackets mark the target-level Wilcoxon signed-rank test of scaling against default across the 13 targets, and the horizontal lines mark TM = 0.85 (dotted) and TM = 0.90 (dashed).

### 2.5 Dissecting the scaling effect

The improvement from scaling grew with the magnitude of the conformational change and was largest on the harder targets (Fig. 4A), and it held at both three and ten recycles in each predictor (Fig. 4B). To test whether scaling acts as a structured modulation of the pair representation or as a generic stochastic perturbation, we added Gaussian noise to the pair representation with a variance matched to the change that scaling induces, and compared the resulting ensembles with those from multiplicative scaling (Fig. 4C, D). In AlphaFold 3 the two operations behaved very differently. Multiplicative scaling preserved well-formed structures, whereas the matched additive noise collapsed the prediction into steric clashes and recovered neither reference state. In Boltz-2 the same additive noise was tolerated, leaving the recovery close to that of scaling. The contrast in AlphaFold 3 shows that the gain from scaling is not reproduced by an unstructured perturbation of equal magnitude and is therefore a structured modulation of the pair representation rather than generic noise. That Boltz-2 absorbed both operations indicates that its pair representation is comparatively robust to perturbation.

**Figure 4.**
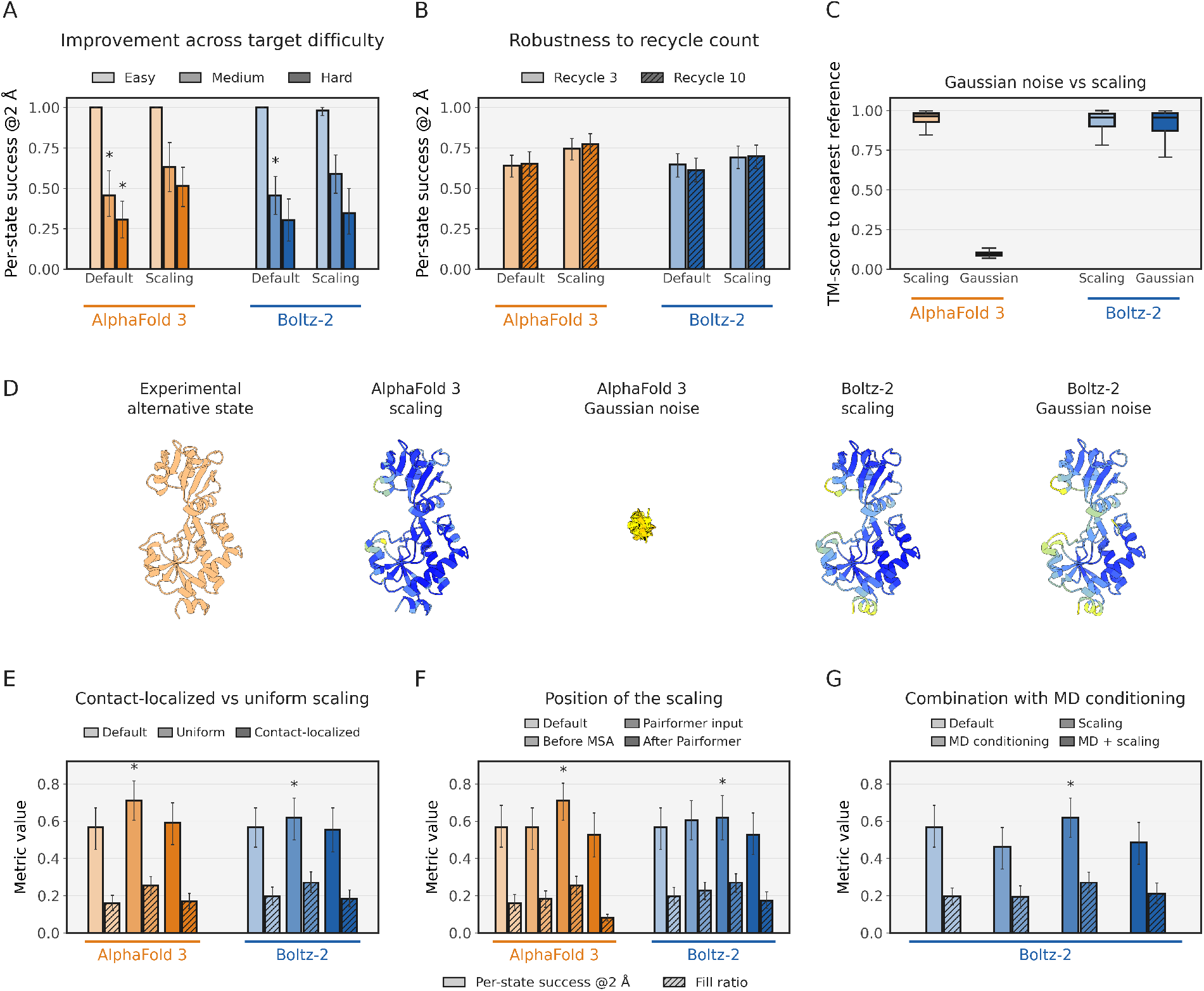
Dissecting pair representation scaling. Throughout, bars are per-target means with 95% bootstrap confidence intervals, and an asterisk marks a variant that significantly improves on default inference within that predictor (paired Wilcoxon, Holm-corrected, *p <* 0.05). **(A)** Per-state success rate at 2 Å for AlphaFold 3 and Boltz-2, stratified by the magnitude of the conformational change (easy, medium, hard), under default inference and scaling. **(B)** The same success rate at three and ten recycles. **(C)** Multiplicative scaling against additive Gaussian noise of matched variance, as the per-sample TM-score to the nearer reference. **(D)** Predicted structures of Q5F9M1 in both predictors under scaling and under the matched noise, beside the experimental alternative state and coloured by pLDDT. **(E–G)** Ablations on a subset of 38 targets, with solid bars the per-state success rate at 2 Å and hatched bars the fill ratio, every condition compared at 150 models per target (Section 5.2). **(E)** default inference, uniform scaling, and contact-localized scaling restricted to the contacts that move between the two reference states. **(F)** scaling applied before the MSA module, at the Pairformer input, or after the Pairformer. Panels E and F show both predictors. **(G)** scaling combined with the MD-conditioned mode of Boltz-2, which is Boltz-2 only because that mode exists only there.

We next asked where in the trunk the scaling has to act, comparing it applied before the MSA module’s outer-product mean, at the input to the Pairformer, and after the Pairformer on a subset of 38 targets composed of the OC23 and IOMemP sets (Fig. 4F). The position mattered (Cochran’s Q across the three, *p* = 0.0001 in AlphaFold 3 and *p* = 0.03 in Boltz-2). Only scaling at the input to the Pairformer, the position used throughout, improved on default inference. Scaling before the MSA module’s outer-product mean did not broaden recovery and left the per-state success rate at the default level, because the outer-product mean then writes the alignment into the pair representation and absorbs the perturbation. Scaling after the Pairformer, directly on the representation that conditions the structure module, instead narrowed the ensemble, halving the fill ratio in AlphaFold 3 and lowering it in Boltz-2 (*p <* 0.001 and *p* = 0.02), so a static change to the structure-module conditioning degrades recovery. The effective handle is therefore the assembled pair representation at the input to the Pairformer, which the outer-product mean builds and the Pairformer then refines before it conditions the structure module. Consistent with this, the response of individual targets to *β* was not monotonic, and across the 86 targets none showed a strictly monotonic change in the recovered TM-score as *β* was varied (Section S5). The benefit of the method comes from sampling across the range of *β* values rather than from a monotonic dependence of structure on *β*.

We then asked whether the effect is carried by the specific contacts that distinguish the two states or by the pair representation as a whole. Scaling restricted to the residue pairs that move between the two reference states, an oracle localization to the contacts whose C*α*–C*α* distance changes by more than 4 Å, was no better than default inference and did not reproduce the gain of uniform scaling in either predictor (Fig. 4E). The benefit therefore comes from the overall magnitude of the pair representation rather than from a spatial pattern that targets the moving contacts.

A final baseline is Boltz-2’s own mechanism for conformational diversity, its MD-conditioned mode. This mode did not by itself broaden recovery beyond default inference, and combining it with scaling did not improve on scaling alone (Fig. 4G). It is built for local dynamics, trained on molecular-dynamics trajectories and reproducing residue-level fluctuations (RMSF) better than dedicated ensemble generators in the Boltz-2 evaluation [19]. Those trajectories stay within a single conformational basin, so the conditioning adds local flexibility, not the large-scale transition between distinct deposited states that this benchmark requires.

#### 2.5.1 Internal representations under scaling

To localise the effect, we read out the internal representations under the full scaling sweep without rescaling them further, taking the post-Pairformer pair representation that conditions the diffusion module and the predicted distance distribution (distogram), which is a linear projection of that same pair representation [17] (Fig. 5). At change-contact pairs, those whose C*β* distance differs by more than 4 Å between the two reference structures, scaling moved the distogram toward the alternative-state geometry in two respects. The fraction of predicted distance mass on the alternative-state side rose more than twofold at the strongest negative *β* in AlphaFold 3, and the distribution broadened (Fig. 5A, B). Both effects appeared in either scaling direction but were larger at negative *β*, and the same quantities moved in the same direction in Boltz-2, only weakly.

**Figure 5.**
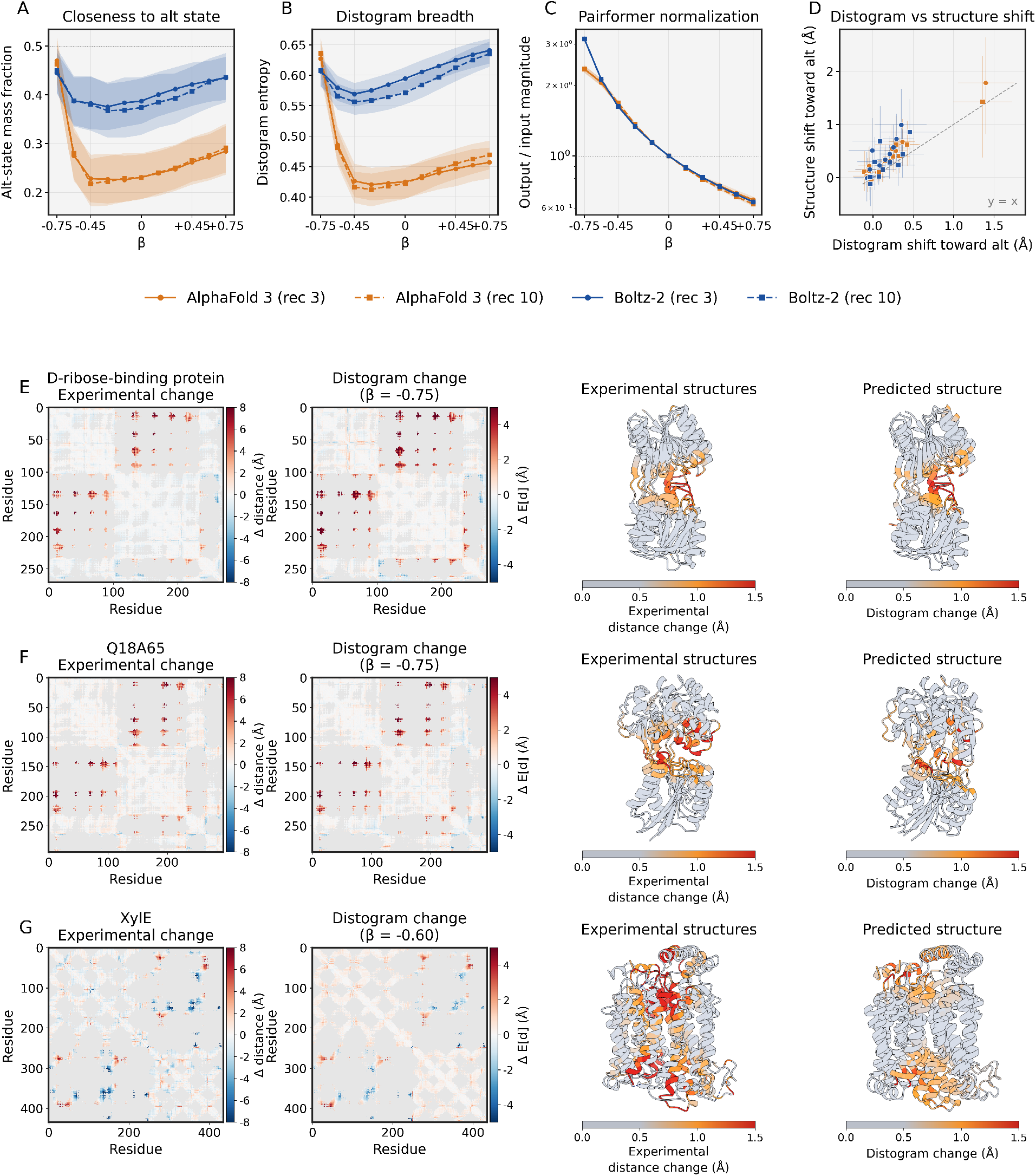
Internal-representation effect of pair representation scaling. **(A)** Distogram mass fraction on the alternative-state side at change-contact pairs versus *β*. **(B)** Distogram breadth (normalized entropy) at change-contact pairs versus *β*. **(C)** Post-Pairformer pair representation magnitude divided by (1 + *β*) versus *β*. A ratio of one is pass-through. **(D)** Per-*β* distogram shift against generated-structure shift toward the alternative state. The dashed line is *y* = *x*. In A–D colour is the predictor and line style the recycle count (solid three, dashed ten), aggregated over the 86 targets with 95% bootstrap bands. **(E–G)** Three AlphaFold 3 examples, the D-ribose-binding protein (E), Q18A65 (F), and XylE (G). Each row shows the experimental distance change (alternative minus dominant state, C*α* distances), the distogram change at *β*^*^, the two experimental references coloured by per-residue distance change, and the two predicted conformers that best match the two reference states, coloured by per-residue distogram change. Both the experimental change and the distogram change are oriented from the dominant toward the alternative state.

The Pairformer attenuates amplified inputs and amplifies shrunken inputs without eliminating the perturbation, as captured by the ratio between the post-Pairformer pair representation magnitude and the input scaling factor (1 + *β*), which sits below unity for *β >* 0 and above unity for *β <* 0 in both predictors (Fig. 5C). The matched additive Gaussian noise had a qualitatively different signature. In AlphaFold 3 the structure module did not produce a folded prediction, whereas in Boltz-2 the noise was largely absorbed (Fig. 4D).

The pair representation enters the structure module’s attention as a bias, so a change in it is carried directly into structure generation. Consistent with this, the per-target distogram shift and the per-target shift of the generated structure, both measured toward the alternative state over the same sweep, were positively correlated, more strongly in AlphaFold 3 than in Boltz-2 (Fig. 5D). To test whether this change is directed toward the experimentally observed alternative state rather than an arbitrary rearrangement, we compared, per target, the distogram change under scaling with the distance change between the two reference structures over the residue pairs within distogram range. The two were positively correlated on nearly all AlphaFold 3 targets with a recovery improvement, with a median Pearson *r* = 0.56 across the 35 such targets and a positive value for 34 of them (Fig. 5E–G). In Q18A65 the residue pairs whose distogram changes are the pairs that differ between the two experimental states, and the generated structure reorganizes at those residues (Fig. 5F).

#### 2.5.2 Distogram bimodality

To resolve whether these distogram shifts reflect a reweighting of pre-existing bimodal distance distributions or the creation of new bimodal hypotheses, we applied the peak-finding protocol of Li et al. [27] to every change-contact pair across all 86 targets and 11 *β* values (Section 5.7). At default inference, about one in ten AlphaFold 3 change-contact distograms were already bimodal, and a majority of those bimodal distributions had peaks at the two reference distances (Fig. 6A–B), whereas Boltz-2 had both a lower baseline and weaker alignment of peaks with the reference distances. These default-inference levels and the alignment of bimodal peaks with the reference distances were similar at three and ten recycles in each predictor (Fig. 6A, B). Under scaling, the bimodal fraction rose significantly in both predictors at both recycle counts (paired Wilcoxon *p <* 10*^−^*^11^ in all four conditions), with the newly bimodal peaks again aligned with the reference distances at a comparable rate to the baseline. The protocol resolves five distinct per-pair response patterns (Fig. 6C). (i) Pairs already bimodal at *β* = 0 whose two peaks track the two reference distances and persist across the sweep. (ii) Pairs unimodal at *β* = 0 that acquire a second peak under scaling, with the new mode landing at the alternative-state distance. (iii) Pairs that stay unimodal but whose single peak shifts continuously between the two reference distances as *β* varies. (iv) Pairs for which scaling leaves the distogram unchanged. (v) Pairs that turn bimodal under scaling at distances matching neither reference, a spurious split rather than the alternative state. About half of the bimodal change-contact pairs fall in this last category in AlphaFold 3 and about three quarters in Boltz-2, so the matched fraction in panel B, rather than the bimodal fraction in panel A, is the measure of whether scaling encodes the alternative state. Per-target histograms for all 86 targets are shown in Figures S11–S16.

**Figure 6.**
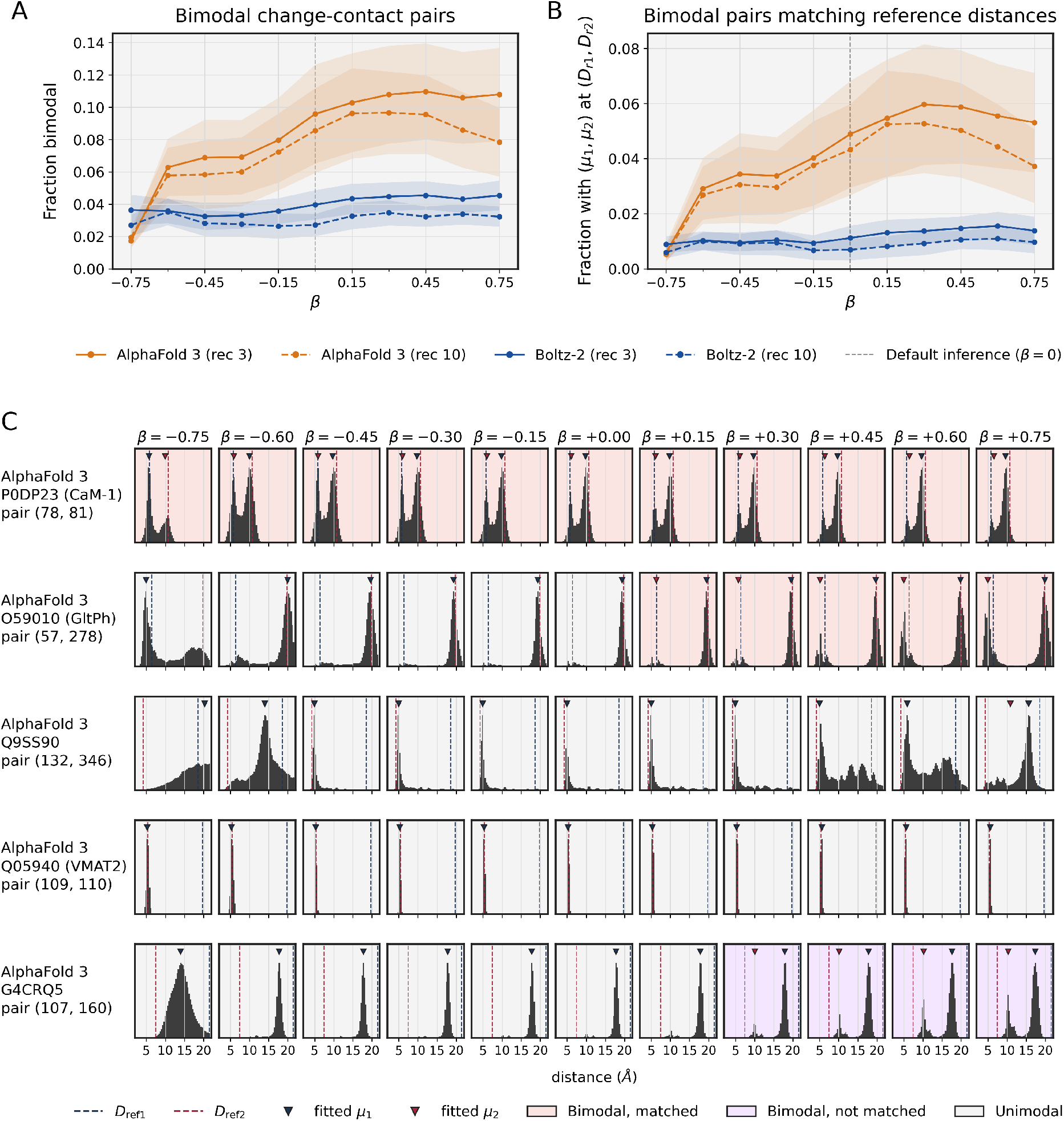
Distogram bimodality under pair representation scaling. **(A)** Fraction of change-contact pairs classified as bimodal versus *β*, for AlphaFold 3 and Boltz-2 at three and ten recycles. Colour is the predictor and line style the recycle count. Lines are target means over 86 targets with 95% bootstrap bands. **(B)** Fraction of change-contact pairs whose fitted bimodal peaks lie within 1.5 Å of the two reference C*β*–C*β* distances, for the same conditions. **(C)** Five representative AlphaFold 3 change-contact pairs at three recycles, one for each response mode described in the text, namely P0DP23 (78, 81), O59010/GltPh (57, 278), Q9SS90 (132, 346), Q05940/VMAT2 (109, 110), and G4CRQ5 (107, 160). Dashed vertical lines mark the two reference distances and triangles the fitted peak positions. The panel background marks the classification, bimodal with both peaks within 1.5 Å of the reference distances (pink), bimodal but matching neither (purple), or unimodal (grey).

#### 2.5.3 Denoising trajectories

The analyses above place the effect of scaling in the trunk, upstream of the diffusion module. To follow how the diffusion module realises this conditioning, we recorded the denoised-structure estimate at each of the 200 AlphaFold 3 denoising steps and measured, at every step, whether it was closer to the dominant or to the alternative reference, over 50 trajectories each under default inference and scaling for two targets (Fig. 7). For both FlgA (P40131) and the post-cutoff antitermination protein Q (P03047), every default trajectory stayed closer to the dominant reference at every step and none reached the alternative side. Under scaling the early estimates are still forming, and once the backbone resolves the trajectory lies on the alternative side, 40 of 50 for FlgA and all 50 for protein Q. The denoised structures overlaid at the first and last step show the backbone displacement toward the alternative state that accompanies this shift (Fig. 7).

**Figure 7.**
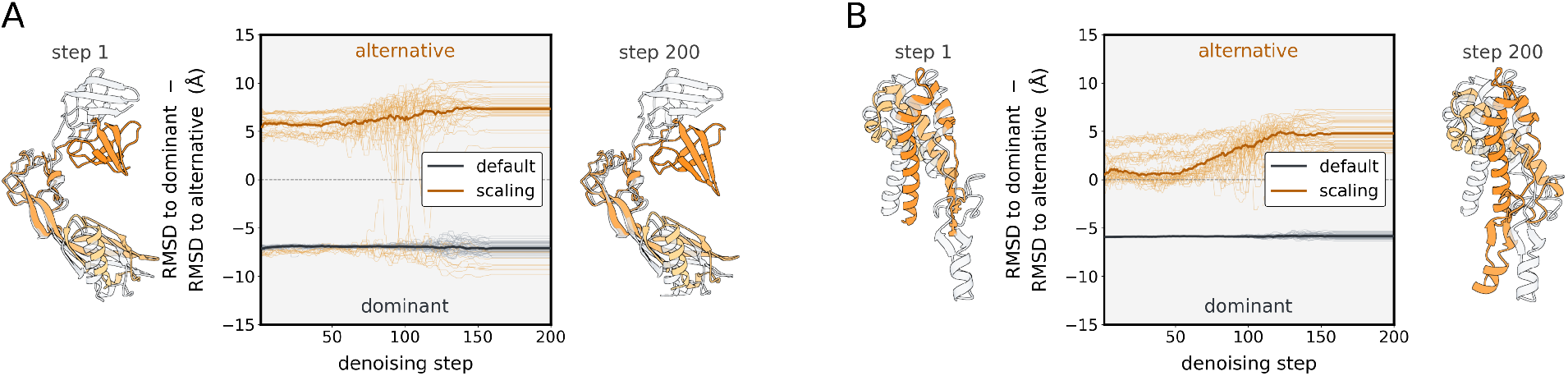
Pair representation scaling sets the sampled state within the denoising trajectory. **(A)** FlgA (P40131) and **(B)** antitermination protein Q (P03047). For 50 trajectories each under default (grey) and scaling (orange), the C*α*-RMSD of the denoised-structure estimate to the dominant reference minus its RMSD to the alternative reference at each denoising step. Medians are bold, and values above zero lie closer to the alternative state. Flanking structures overlay a representative dominant (grey) and alternative (orange) trajectory at the first and last step. The first-step overlays are early denoised estimates and can be only partially formed.

### 2.6 Sequence-only inference

The benchmark above used the multiple sequence alignment that each predictor would normally receive. To test whether pair representation scaling depends on coevolutionary input, we repeated the evaluation with the alignment removed, so that each prediction was driven by the single query sequence (Fig. 8). Removing the alignment reduced accuracy sharply for both predictors, and the coverage fell below 0.10 for every method. Even so, scaling retained a measurable benefit. In AlphaFold 3 both metrics shifted in the same direction under scaling (Fig. 8A). A per-target comparison confirmed this direction (Fig. 8B). Of the 86 AlphaFold 3 targets, scaling improved the worst-case minimum RMSD by at least 1 Å on more targets than it degraded, so the operation helped more targets than it harmed even without an alignment. Scaling therefore persisted in the absence of a coevolutionary signal.

**Figure 8.**
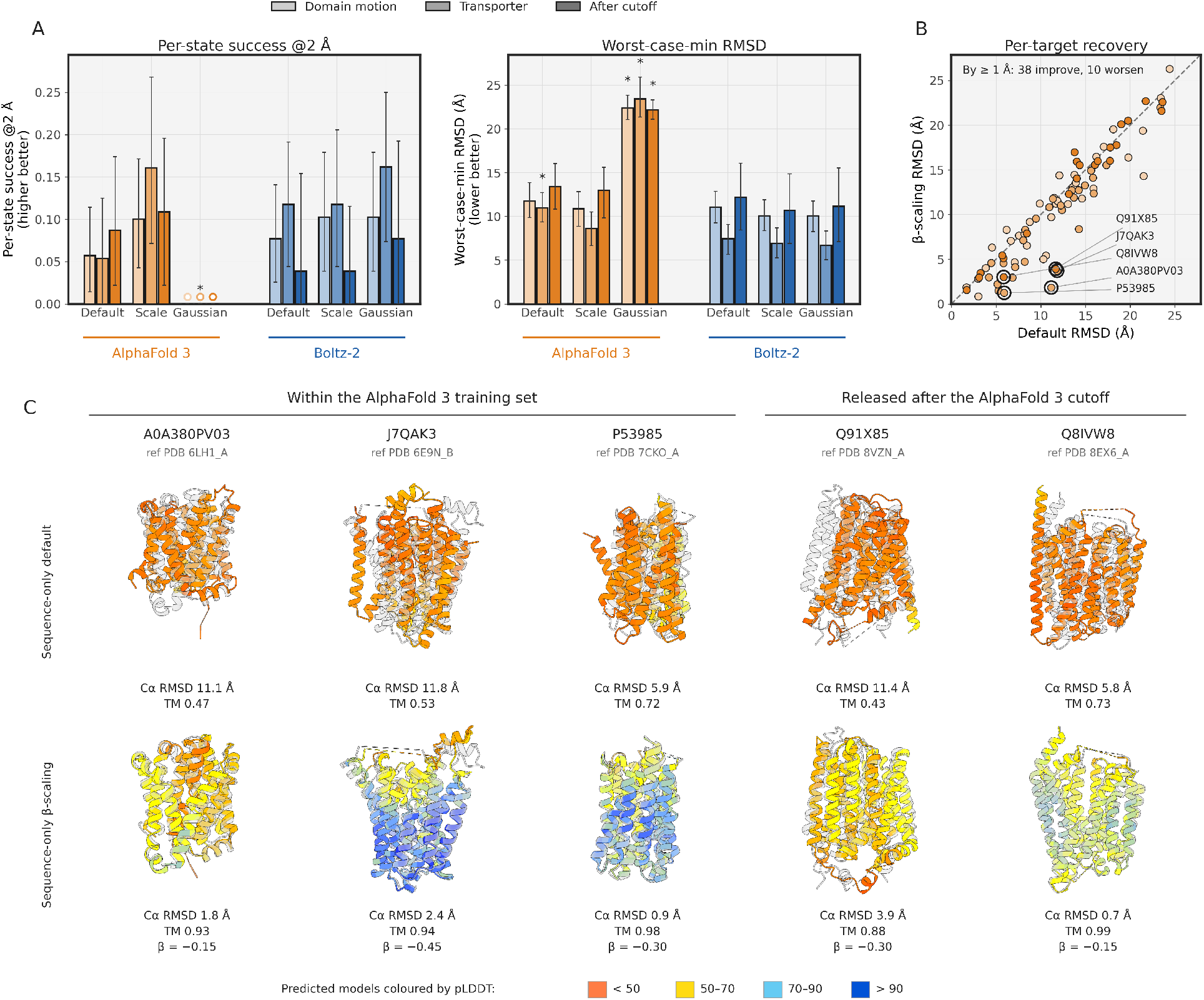
Sequence-only inference. **(A)** Per-state success rate and worst-case minimum RMSD without a multiple sequence alignment, by group and predictor, under default inference, scaling, and the matched Gaussian-noise control. An asterisk marks a method that scaling significantly outperforms (paired Wilcoxon, Holm-corrected, *p <* 0.05). **(B)** Per-target worst-case minimum RMSD, default against scaling, AlphaFold 3 only. The five case-study targets of panel C are ringed. **(C)** Five membrane transporters under sequence-only AlphaFold 3 inference. The best default and best negative-*β* scaling models are coloured by pLDDT and overlaid on the experimental reference in grey, with C*α* RMSD and TM-score below each. Targets are labelled by UniProt accession, the first three in-training and the last two post-cutoff.

The per-target gain was concentrated rather than uniform, and for some targets it was large (Fig. 8C). For five membrane transporters, sequence-only default inference in AlphaFold 3 produced a misfolded model far from the experimental reference whereas negative-*β* scaling recovered a near-native structure (Fig. 8C, with per-target RMSD and TM-score values reported below each prediction). Two of these five targets have reference structures released after the AlphaFold 3 training cutoff, so the rescue extends to structures the model had not seen during training.

Negative *β* was the more helpful direction both with and without an alignment, and the sequence-only rescues came consistently from negative *β* (Fig. 8). Which of the two states a given *β* surfaces, however, stays target-specific (Section S3). The matched Gaussian-noise control, run here without an alignment, again remained catastrophic for AlphaFold 3 and was tolerated by Boltz-2.

## 3 Discussion

A single scalar multiplication of the latent pair representation broadens the conformational ensembles produced by two diffusion-based structure predictors, and the controls identify what kind of intervention this is. Matched additive noise does not reproduce the gain, and scaling restricted to the contacts that move between the two states is no better than uniform scaling, so the effect is a structured, global change in the magnitude of the pair representation rather than added stochasticity or a spatial pattern. The perturbation also has to enter at the Pairformer input and be re-processed by the trunk, since scaling before the MSA module does not broaden the ensemble and rescaling the structure-module conditioning directly degrades it. The effective handle is therefore a global modulation of the pair representation upstream of the Pairformer, which is why the same operation transfers unchanged between AlphaFold 3 and Boltz-2. Because the operation changes the magnitude of a single internal representation without touching the input alignment, the same effect can be measured inside the network, retained without an alignment, and combined with alignment-based sampling.

The distogram analysis shows how this modulation reaches structure. Already at default inference a fraction of AlphaFold 3 change-contact distograms are bimodal with modes at both reference distances, so the model encodes the two-state distribution in the pair representation before any intervention, consistent with the finding that AlphaFold distograms capture a hinge motion that the single predicted structure does not [28]. Recent probing work finds biophysical features linearly accessible in the AlphaFold 3 pair latent [29], and in ESMFold counterfactual interventions localize distance and contact information to the pairwise representation and steer the predicted structure through it [30, 31]. Under scaling more pairs become bimodal and others shift their single peak between the two references, so the operation both reweights existing bimodal hypotheses and extends the reach to new ones, and because these changes track the experimental distance change between the two states the rearrangement it induces is directed toward the observed alternative state rather than arbitrary. The diffusion module carries this conditioning into the sampled structure within the denoising trajectory, which under scaling resolves to the alternative state while the default trajectory stays in the dominant state at every step (Fig. 7). The same picture is weaker in Boltz-2, where the matched noise is tolerated rather than catastrophic at the same recycles.

The effect was larger on AlphaFold 3 than on Boltz-2, consistent with this comparative robustness of the Boltz-2 pair representation. The predictors differ in architecture, training data, and training objective together, so the source of that robustness is not attributable to any single factor. Consistent with this, scaling alone is sufficient in AlphaFold 3, whereas in Boltz-2 the harder state is recovered only when scaling is combined with alignment subsampling, a combination of an internal-representation perturbation with an input one that the MD-conditioned mode does not provide. Negative *β* was the more effective direction for most targets both with and without an alignment, and more uniformly so without it, so this directional preference depends on the strength of the coevolutionary signal the pair representation carries. Related inference-time methods reach alternative conformations through sequence association in AlphaFold 2 [32] or engineered alternate-frame constructs in AlphaFold 3 [33].

Because the distogram is a linear projection of the pair representation, a natural extension is to guide the distogram toward a known alternative-state distance pattern, building on the use of AlphaFold distograms as structural constraints [34, 27]. Coupling structure generation with physical or molecular-dynamics signals, through force-guided sampling [35] or smoothed molecular dynamics [36], is a further direction.

These gains are partial and target-dependent rather than uniform. Scaling widens the ensemble toward the alternative state. Without a reference structure it does not identify which sampled model is the alternative state, because the helpful direction of *β* is target-specific. The alternative state also occupies a small part of the ensemble where it is recovered at all, a median of 13 of 250 models in the AlphaFold 3 domain-motion group, and the gain is accompanied by a loss of occupancy of the dominant state (Section S2). Some targets are not driven to the alternative state at any value of *β*, and in Boltz-2 only the fill-ratio gain over default inference reached significance. The distogram account is partial in the same way, as the bimodality and the mode shifts under scaling appear in a fraction of the change-contact pairs and targets rather than across all of them. Because the Pairformer pushes the imposed magnitude change back toward unity without eliminating it (Fig. 5C), the effect is carried by the part of the change that survives the trunk, and why that surviving signal is directed toward the alternative state remains open. The sequence-only benefit operates where the base prediction is already poor, so it rescues isolated cases rather than providing a general route. The intermediate-state analysis tests structural similarity to deposited structures between the two states rather than kinetic intermediacy of the transition. The after-cutoff group separates targets by the release date of their reference structures, which controls for exposure to the deposited coordinates alone, so a homolog of a post-cutoff target may still have been present in the training set. A split on sequence or structure similarity to the training set would be a stronger control. The benchmark consists of two-state proteins, so behaviour on intrinsically disordered regions, multi-state systems, complexes, and ligand-induced rearrangements [37] remains to be established. More broadly, the method surfaces alternative states without assigning them equilibrium populations, and how inference-time perturbations relate to the underlying conformational Boltzmann distribution remains an open question [38].

## 4 Conclusion

Pair representation scaling broadens the conformational ensemble of a modern diffusion-based structure predictor through a single global modulation of the pair representation upstream of the Pairformer, applied identically to AlphaFold 3 and Boltz-2. The predicted distance distributions show why, as the two-state distribution is already encoded in the pair representation and scaling reweights and extends it toward the experimentally observed alternative state. The modulation retains a benefit without an alignment and recovers the harder Boltz-2 state in combination with alignment subsampling. The effect is partial and target-dependent, but it comes from a single global scalar, making the method a low-cost and interpretable handle on conformational sampling rather than a complete account of it.

## 5 Methods

### 5.1 Implementation of the scaling operation

#### 5.1.1 Uniform scaling

Both AlphaFold 3 and Boltz-2 process a latent pair tensor *z* ∈ R^N*×*N*×*d^*^P^* in their Pairformer trunk. Pair representation scaling rescales this tensor at the input to the Pairformer at each recycling iteration. For uniform scaling, a single scalar *β* multiplies every residue pair,

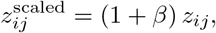

so that *β* = 0 recovers default inference. We swept *β* over {±0.15, ±0.30, ±0.45, ±0.60, ±0.75}, a range chosen to perturb the generated ensemble while avoiding the projection collapse and steric clashes that arise at more extreme values. To locate where the operation must act, we also applied uniform scaling before the MSA module and after the Pairformer, in place of its default position at the Pairformer input.

#### 5.1.2 Contact-localized scaling

To test whether the effect depends on which residue pairs are scaled, we localized the scaling to the contacts that move between the two reference states. For each target we built a binary mask of change-contact pairs, the pairs (*i, j*) whose C*α*–C*α* distance differs by more than 4 Å between the two reference structures (the distogram analysis of Section 5.7 instead uses C*β*), taken over the residues resolved in both states and indexed in the model input-sequence order. Scaling was then applied as 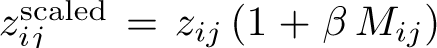with *M*_ij_ the mask, so that only the contacts known to move were rescaled while the rest of the pair representation was left unchanged. Because the mask is built from the deposited reference structures, this is an oracle localization against which uniform scaling can be compared.

#### 5.1.3 Gaussian-noise control

To separate a structured modulation of the pair representation from a generic stochastic perturbation, we used a calibrated Gaussian-noise control. This control adds isotropic Gaussian noise to the pair representation, 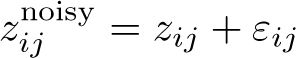 with *ε*_ij_ ∼ N (0*, σ*^2^), and sets the noise variance to the change in element variance that uniform scaling by the same *β* would produce, *σ*^2^ = |(1 + *β*)^2^ − 1| Var(*z*), with Var(*z*) estimated per feature channel. The control therefore matches the magnitude of the perturbation that scaling applies but discards its direction, and it was swept over the same *β* grid.

### 5.2 Datasets

We evaluated 86 targets, each a protein with two experimentally determined conformations, of two structural kinds, all taken from previously published benchmarks. The 86 targets cover 85 proteins. The lipid II flippase MurJ contributes one target to each class, with a different reference pair in each. The individual targets are listed in Table S1. Sequences were taken from UniProt [39] and paired reference structures from the Protein Data Bank [40], and each target was defined at the construct level.

#### 5.2.1 Domain-motion proteins

The 39 domain-motion targets undergo hinge and lid rearrangements. They combine the 22 domain-motion proteins of BioEmu [23] with the 17 proteins of the OC23 set of AFsample2 [9, 41] that BioEmu does not already contain. Each target keeps the reference pair its source benchmark defines.

#### 5.2.2 Membrane transporters

The 47 membrane transporters interconvert between inward-facing and outward-facing states. They combine 15 of the 16 transporters of the IOMemP benchmark [42] with 32 of the 37 membrane proteins in the entropy-guided folding benchmark [13], after removing five duplicates of IOMemP targets. The remaining IOMemP transporter, SPF1, was left out for its length of about 1,090 residues.

#### 5.2.3 Training-cutoff grouping

To test whether the gain depends on the alternative state having been seen during training, we grouped the targets by their reference release date relative to each predictor’s training cutoff. The cutoffs were taken from the model papers, namely 2021-09-30 for AlphaFold 3 [17], 2023-06-01 for Boltz-2 [19], and 2023-11-23 for BioEmu [23], and the release date of every reference structure was verified against the RCSB Protein Data Bank. For each predictor the targets whose later reference was released after its cutoff form an after-cutoff group, holding 23 targets for AlphaFold 3, 13 for Boltz-2, and 10 for BioEmu, and the remaining targets are grouped by structural kind.

#### 5.2.4 Ablation subset

The contact-localized scaling, the alternative scaling positions, and the combination with MD-conditioned inference were each evaluated on a subset of 38 targets, the 23 OC23 domain-motion proteins and the 15 IOMemP transporters. These conditions, together with the Gaussian-noise control, were generated with 150 models per target over the same *β* grid, and the ablation panels compare every condition at that budget, capping default inference and uniform scaling by the rule the benchmark applies at 250 models.

### 5.3 Multiple sequence alignments

Input alignments were taken once from the AlphaFold 3 server [17] and reused unchanged across every method. AlphaFold 3 subsamples the alignment to a fixed depth at each forward pass, and the Boltz-2 runs used a matched depth of 1024 sequences (the AlphaFold 3 server default), while the sequence-only experiments were run with the alignment removed.

### 5.4 Baselines

We compared the scaling operation against default inference and a panel of alternative sampling methods, each generating 250 models per target with the budget split across seeds, settings, and diffusion samples as listed in Table 1. AlphaFold 3 was run locally.

**Table 1.** Sampling methods and settings.

| Method | Setting | Seeds | Settings | Samples | Models |
| --- | --- | --- | --- | --- | --- |
| Default inference | – | 5 | 1 | 50 | 250 |
| MSA subsampling | depth $\in \{2, 4, \dots, 1024\}$ | 5 | 10 | 5 | 250 |
| Random masking | masking rate 0.40 | 5 | 10 | 5 | 250 |
| MSA clustering | $k = 10$ clusters | 5 | 10 | 5 | 250 |
| Inference-time dropout | rate 0.25 | 5 | 1 | 50 | 250 |
| MD-conditioned inference (Boltz-2) | MD method conditioning | 5 | 1 | 50 | 250 |
| Pair representation scaling | $\beta \in \{\pm 0.15, \dots, \pm 0.75\}$ | 5 | 10 | 5 | 250 |
| BioEmu | default settings | 1 | 1 | 250 | 250 |

Three baselines act on the input alignment. MSA subsampling randomly retains a fixed number of the aligned sequences, which weakens the global coevolutionary signal [6]. Random masking selects a random fraction of the MSA columns and replaces each across all sequences with a mask token, weakening the coevolutionary signal, at the 40% rate that AFsample3 found best for AlphaFold 3 [10]. MSA clustering partitions the alignment into *k* = 10 subsets that carry distinct coevolutionary signals, following the AF-Cluster method [11]. We set *k* = 10 to match the ten settings used for the other sweep-based baselines and held it fixed across targets. Whereas the original approach maps the full evolutionary landscape by exhaustive clustering [11], we selected a fixed number of representative clusters with a ranking score *S*_c_ that favours subsets with distinct coevolutionary signals, indicated by a higher internal distance, while retaining sequence depth, indicated by a larger cluster size. Following the AF-Cluster protocol, we removed sequences with an internal gap fraction above 0.25 after trimming terminal gaps, retaining the query sequence regardless of this threshold. From up to 2,500 seed sequences we computed a normalized gap-free edit distance between 0 and 1. We clustered this distance matrix with DBSCAN [43], requiring at least four sequences to form a cluster and choosing, over a neighbourhood radius swept from 0.05 to 0.50, the value that produced the most clusters. The clusters were ranked by

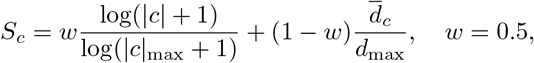

where |*c*| is the cluster size, *d*_c_ its mean internal distance, and *d*_max_ the largest seed distance. For each of the top *k* = 10 clusters we identified the medoid, the sequence with the minimal sum of distances to the other members, and assigned every sequence from the full filtered set to its nearest medoid to form the final ten MSA subsets used for prediction.

Two further baselines act during inference. Inference-time dropout enables dropout at a rate of 0.25 in the Pairformer and MSA stacks during sampling, following AFsample [12], identically in AlphaFold 3 and Boltz-2. Boltz-2 conditions its structure module on the experimental method of the target structure, and its default inference conditions on X-ray diffraction [19]. As a further baseline we evaluated the MD-conditioned mode of Boltz-2, which instead conditions the structure module on molecular dynamics. We also compared against BioEmu, a separate generative model trained to emulate protein equilibrium ensembles [23], run with its default settings and evaluated with the same pipeline as the other methods.

### 5.5 Evaluation metrics

Every generated model was compared against both reference states by TM-score and C*α* RMSD, and each ensemble was summarized with the metrics below.

#### 5.5.1 Per-state recovery

The per-state success rate is the fraction of the two reference states for which the ensemble contains a model within 2 Å C*α* RMSD. The worst-case minimum RMSD is the larger of the two minimum RMSD values to the references and measures recovery of the harder state. The best-minimum TM-score, 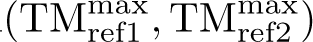, is the TM-score of the harder state at the best-scoring model for each reference.

#### 5.5.2 Fill ratio

Following AFsample2 [9], the fill ratio measures how completely the ensemble covers the path between the two reference states. Each model sits at 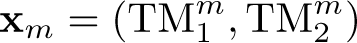, its TM-scores to the two references, which lie at *A* = (1*, τ*) and *B* = (*τ,* 1) with *τ* the reference–reference TM-score, so the segment *AB* is the path between the two states. Each model that improves on both references 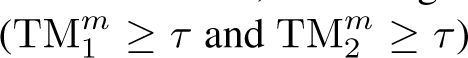 is projected onto *AB* at the fractional position 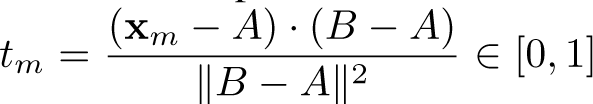 We split the path into *N* = 100 equal bins and count the bins that hold at least one model, *N*_occ_. The fill ratio is their fraction (Equation 1),

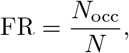

so FR = 1 when the ensemble spans the whole path and falls toward 0 as the models collapse onto a single point.

#### 5.5.3 Statistical testing

All comparisons were paired across targets within each conformational-change class. Pair representation scaling was compared against each baseline with a two-sided paired Wilcoxon signed-rank test on per-target metric values, Holm-corrected within each predictor, class, and metric. Binary recovery across the three scaling positions was compared with Cochran’s Q test followed by McNemar’s exact test. Effect sizes are reported with 95% confidence intervals from 10,000 paired bootstrap resamples.

### 5.6 Recovery of deposited intermediate structures

For each of the 39 domain-motion targets we retrieved every PDB entry cross-referenced from the target’s UniProt record, removed the two reference structures, and downloaded the remaining entries from the RCSB. We computed the TM-score from each candidate to both references, and a candidate qualified as an intermediate when both TM-scores fell between 0.70 and 0.95, the upper bound excluding near-duplicates of either reference state and the lower bound enforcing the same fold. The paired direction, significance, and effect-size magnitude were unchanged when the lower bound was varied between 0.65 and 0.75 and the upper bound between 0.93 and 0.97. For each intermediate and condition we drew 100 predictions without replacement from the corresponding ensemble at a fixed seed, computed the TM-score from the intermediate to each, and kept the maximum. Best TM-score was compared between default and scaling within each predictor with a one-sided Wilcoxon signed-rank test on the 75 paired values, the direction set by the hypothesis that scaling brings the ensemble closer to the intermediates, and threshold counts with the McNemar exact test. Because the 75 intermediates come from 13 targets, we confirmed the comparison at the target level by averaging the improvement within each target and testing across the 13 targets.

### 5.7 Distogram bimodality analysis

We applied the peak-finding protocol of Li et al. [27] to test whether the predicted distance distribution at the input to the structure module is bimodal at change-contact residue pairs. For each prediction we read out the distogram probabilities over the 64 distance bins spanning 2 to 22 Å, treating the final bin as long-distance overflow. The two highest-probability local maxima were taken as bimodality candidates when their combined probability exceeded 0.1 and the secondary peak retained at least a tenth of the primary peak. A sum of two Gaussians was fit to these candidates by least squares, rejecting fits whose fitted standard deviation reached 3 Å or whose mean fell beyond the distogram range. A pair was classified as bimodal when the fitted means were separated by at least 1.5 Å and the probability at the saddle between them dropped at least 5% below the secondary peak, and as matching the two reference states when the fitted means lay within 1.5 Å of the two reference C*β*–C*β* distances in either ordering.

Change-contact pairs were the residue pairs whose C*β*–C*β* distance differed by more than 4 Å between the two reference structures with both reference distances inside the distogram’s representable range, using C*β* for non-glycine residues and C*α* for glycine. The predictor input sequence was aligned to each reference independently and the shared aligned positions were used to index the distogram. Classification used the protocol’s published default thresholds, a secondary-peak probability ratio of 0.10 and a saddle-point prominence of 0.05.

### 5.8 Denoising-trajectory analysis

To follow how the diffusion module realises the scaled conditioning, we instrumented the AlphaFold 3 sampler to record the denoised-structure estimate at each of the 200 denoising steps, for FlgA (P40131) and the post-cutoff antitermination protein Q (P03047). For each target we generated 50 trajectories each under default inference and under scaling at the per-target *β*^&^ (−0.75 and −0.60), as five seeds of ten samples. The per-step estimate lives in a random global frame because of the diffusion augmentation, so each was superimposed on the two references over the static core, the half of the aligned residues whose position differs least between the two reference states, and assigned to a state by the sign of its C*α*-RMSD to the dominant reference minus its RMSD to the alternative reference. The overlaid structures show one representative dominant and one alternative trajectory, oriented so the displacement between the two states lies in the image plane.

### 5.9 Software and hardware

Structures were generated with AlphaFold 3 version 3.0.1 [17], Boltz-2 version 2.2.1 [19], and BioEmu version 1.3.1 [23], each at its released default inference settings, with the recycle count of the two diffusion-based predictors fixed to three (Table 2). TM-scores and C*α* RMSD values were computed with US-align [44], and structures were rendered with ChimeraX [45]. Predictions were run on NVIDIA RTX A6000 and H100 GPUs.

**Table 2.** Inference settings for the two diffusion-based structure predictors. All values are the released defaults except the recycle count, set to three for both.

| Setting | AlphaFold 3 | Boltz-2 |
| --- | --- | --- |
| Version | 3.0.1 | 2.2.1 |
| Training data cutoff | 2021-09-30 | 2023-06-01 |
| Recycling iterations | 3 | 3 |
| Diffusion denoising steps | 200 | 200 |
| Diffusion noise scale ($\lambda$) | 1.003 | 1.003 |
| Diffusion step scale ($\eta$) | 1.5 | 1.5 |
| Noise-inflation factor ($\gamma_0$) | 0.8 | 0.8 |
| MSA depth | 1024 | 1024 |
| Input templates | none | none |

### 5.10 Data and Software Availability

The implementation is available in a public repository at https://github.com/suzuki-2001/pair-representation-scaling.

## Computational resources

This research used computational resources of Pegasus at the Center for Computational Sciences, University of Tsukuba, through the Multidisciplinary Cooperative Research Program, and of SQUID [46] at the D3 Center, The University of Osaka, through the Project for Nurturing Student Competing with the World.

## Ethical Statement

This study did not involve human participants or animals, and therefore ethical approval was not required.

## Author Contributions

S.S conceived and carried out the study, performed the analyses, and wrote the manuscript. T.A supervised the research and provided critical feedback on the manuscript. Both authors read and approved the final version of the manuscript.

## Supporting Information

Table of the 86 benchmark targets with UniProt accessions, reference structures and chains, release dates, and per-predictor difficulty; distribution across targets of the ensemble occupancy of each reference state, and best-minimum TM-score by conformational-change group; per-state success as a function of ensemble size for AlphaFold 3 and Boltz-2; per-target Spearman correlation between *β* and the conformational state; reference-free two-model selection curves for five selection rules; distogram–experiment agreement for every AlphaFold 3 target with a recovery improvement; per-target TM-score scatter to both reference states for all 86 benchmark targets; per-target AlphaFold 3 distogram bimodality across the *β* sweep for all 86 benchmark targets (PDF)

## Supporting Information

### S1 Benchmark targets

Table S1 lists the 86 benchmark targets. The source benchmarks are the domain-motion set of BioEmu [23], OC23 [9], IOMemP [42], and the entropy-guided folding membrane set (EGF) [13]. The BioEmu and OC23 targets form the domain-motion class and the IOMemP and EGF targets the transporter class. Six targets, Q5F9M1, Q18A65, A2RJ53, A0A075Q0W3, P33284 and P71447, belong to both the BioEmu and the OC23 set. They are listed once, under OC23, and their reference pairs are the ones BioEmu specifies, which is why the Methods count them among the 22 BioEmu proteins and take 17 further proteins from OC23. Difficulty is defined per predictor on that predictor’s own default ensemble, binning its worst-case minimum C*α* RMSD as easy (*<* 2 Å), medium (2–4 Å), or hard (≥ 4 Å), so a target can be easy for one predictor and hard for the other. The UniProt accessions, protein names and release dates were read from the RCSB Protein Data Bank.

**Table S1.**
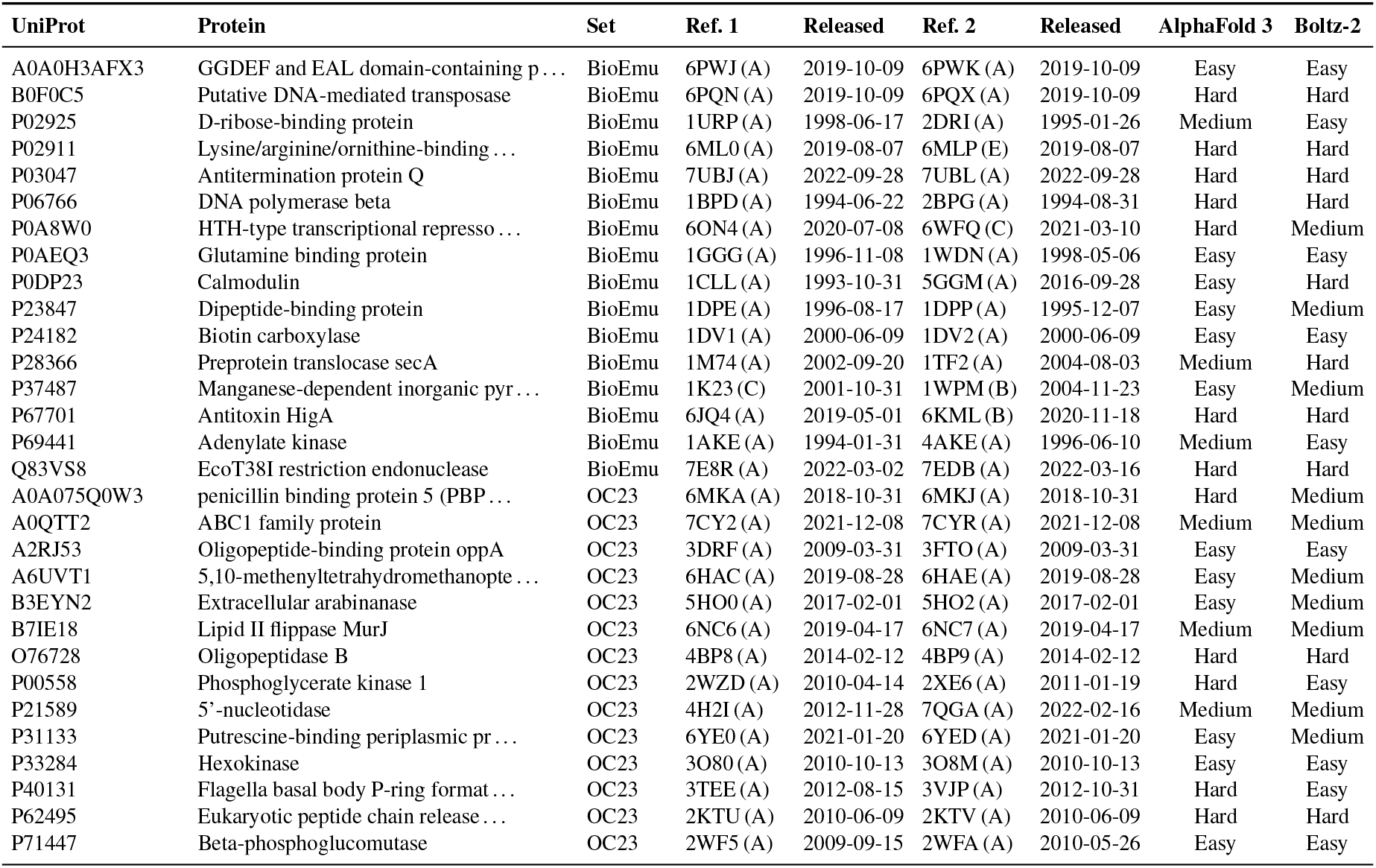
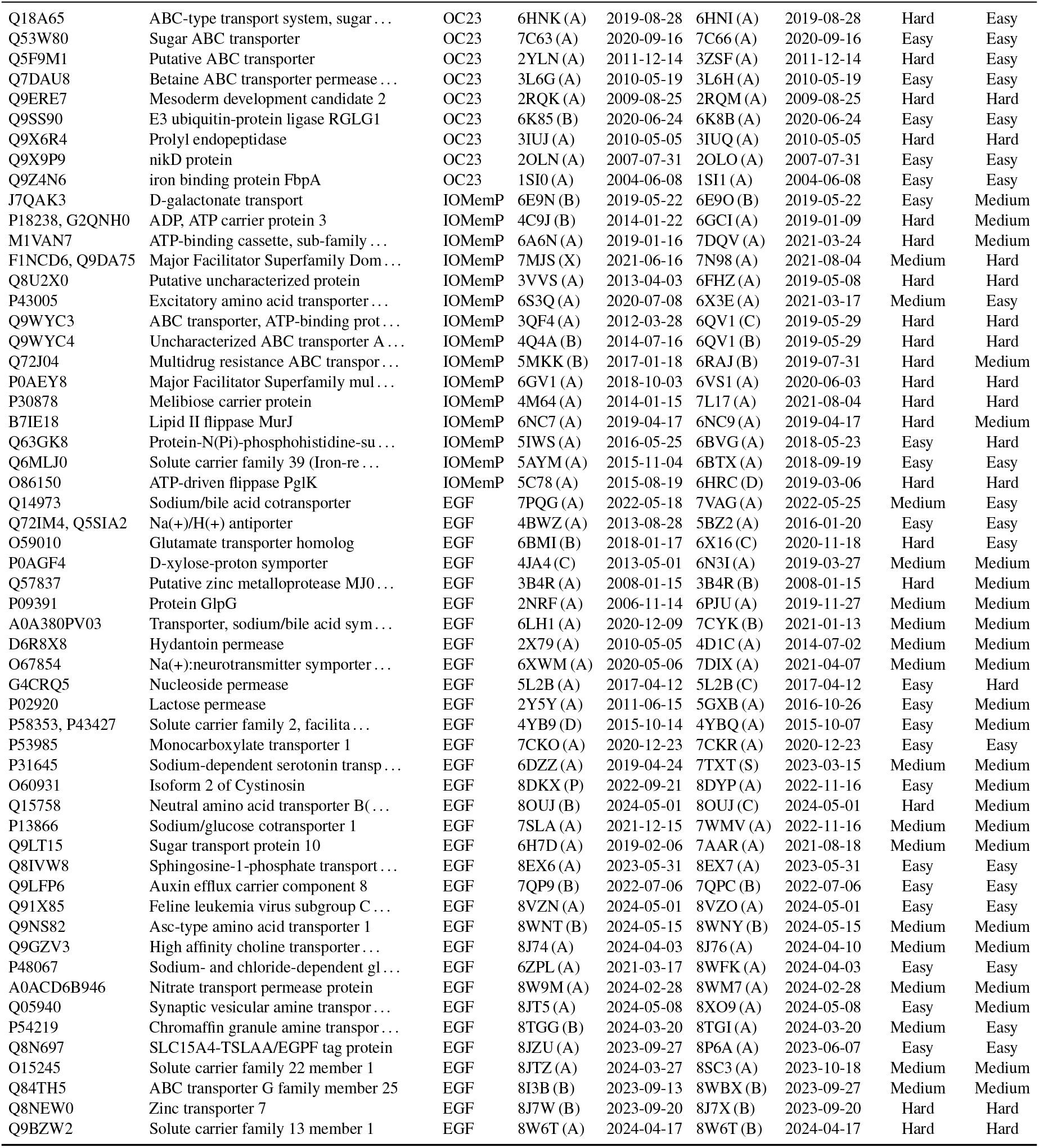
The 86 benchmark targets. UniProt accession, protein, source benchmark, the two reference structures with their author chains and Protein Data Bank release dates, and the difficulty assignment for each predictor. For 4 targets the two references come from different organisms or strains of the same protein, and both accessions are listed.

### S2 Ensemble occupancy of the two states

Occupancy of a state is the fraction of the 250-model ensemble within 2 Å C*α* RMSD of that reference, measured here for every target and method (Fig. S1).

Occupancy of the harder state is low throughout. Under default inference the ensemble never reaches that state for 63% of the AlphaFold 3 targets and 66% of the Boltz-2 targets, and scaling lowers those fractions to 44% and 55% (Fig. S1A,B). Where the state is reached the occupancy stays small, with the per-target median across the domain-motion group rising from 0 to 0.052 in AlphaFold 3, or 13 of 250 models. Scaling raises the harder-state occupancy over default inference in all three AlphaFold 3 groups and in the Boltz-2 domain-motion and after-cutoff groups (paired Wilcoxon signed-rank test, *p* ≤ 0.046), but not in the Boltz-2 transporter group (*p* = 0.73).

The gain comes with a shift away from the dominant state (Fig. S1C). Following the reference that the default ensemble covers better, the median occupancy of that state falls from 0.97 to 0.74 in AlphaFold 3 and from 0.86 to 0.73 in Boltz-2, and it decreases for 56 of the 72 and 56 of the 74 targets whose default ensemble reaches it at all. Scaling therefore redistributes ensemble mass between the two states rather than adding coverage of the alternative state on top of the dominant one.

The best-minimum TM-score rises under scaling in every group of both predictors, by 0.001 to 0.040, but the 95% bootstrap confidence intervals of the paired conditions overlap in all six, so it separates the conditions less sharply than the three metrics of Figure 1 (Fig. S1D).

**Figure S1.**
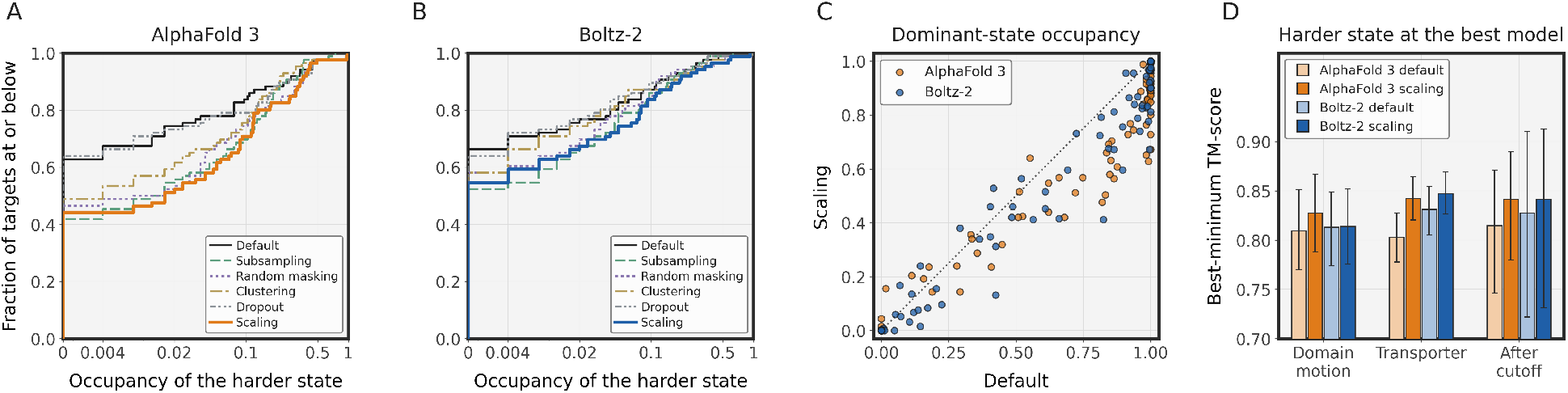
Ensemble occupancy of the two reference states. Occupancy is the fraction of the 250-model ensemble within 2 Å C*α* RMSD of a reference state. **(A, B)** Distribution across targets of the occupancy of the harder state, as an empirical cumulative distribution, for AlphaFold 3 (A) and Boltz-2 (B). The value at zero occupancy is the fraction of targets whose ensemble never reaches that state. Scaling is drawn in the predictor colour and the baselines in the lighter tint. **(C)** Occupancy of the dominant state per target, default inference against scaling. The dominant state is the reference that the default ensemble covers better, and the same reference is followed into the scaled ensemble. The dotted line is the identity. **(D)** Best-minimum TM-score by conformational-change group, as target means with 95% bootstrap confidence intervals.

### S3 Sample efficiency and reference-free selection

We first asked whether the scaling parameter itself carries information about the conformational state (Fig. S3A). Within each target we measured the Spearman correlation between *β* and the difference in TM-score to the two reference structures across the scaled models. The correlation was moderate and significant for most targets, more strongly in AlphaFold 3 than in Boltz-2 (Fig. S3A). The sign of the effective direction was target-specific, positive for roughly half the targets and negative for the rest, so no single direction of *β* maps onto a given conformational state across targets.

The broadening did not come from drawing more models. We resampled each method’s ensemble at random, from a single model up to the full ensemble, and recomputed two-state per-state success at each size (Fig. S2A,B). Scaling reached the per-state coverage of the full default ensemble from a fraction of it, about a tenth in AlphaFold 3 and under half in Boltz-2, and stayed above every baseline across the whole range. The gain over default inference is therefore a property of the per-sample distribution rather than of the number of models drawn.

We then tested whether two models covering both conformational states could be selected from a scaled ensemble without a reference structure (Fig. S3B,C). A rule returned two models, scored by the area under a success curve, where success at a TM-score threshold means that the two models include one within that threshold of each reference structure. The per-target correlation between *β* and the state did not translate into a selection signal. Sign-based selection matched a random baseline for both predictors. Mean pLDDT peaked near default inference and declined with the magnitude of *β*, so the most confident models concentrated on the dominant state and selection by the two highest-pLDDT models fell below the random baseline. Structural clustering of the ensemble’s secondary-structure-region C*α* distance maps, returning the two cluster medoids, recovered both states more often than chance and approached the oracle ceiling for both predictors (Fig. S3B,C). The scaled ensemble therefore holds both conformational states in a form that an unsupervised structural criterion can separate, whereas the predictor’s confidence is biased toward the dominant state and the *β* sign, though correlated with the state, is too target-specific to act on without a reference.

#### Selection rules

The success curve underlying each rule varies the TM-score threshold from 0.5 to 1.0, and a target counts as recovered at a threshold when the two returned models include one within it of the first reference structure and one within it of the second. Sign-based selection draws one model at random from the models generated with *β <* 0 and one from those with *β >* 0. Confidence selection returns the two models of highest mean C*α* pLDDT. Structural clustering partitions the ensemble into two groups by *k*-means with *k* = 2 on the C*α* distance maps restricted to the consensus secondary-structure residues, those assigned to a helix or strand by DSSP in at least half of the models, and returns the medoid of each group. The random baseline draws two models uniformly, and the oracle is the best two-model coverage attainable from the ensemble with knowledge of the reference structures.

**Figure S2.**
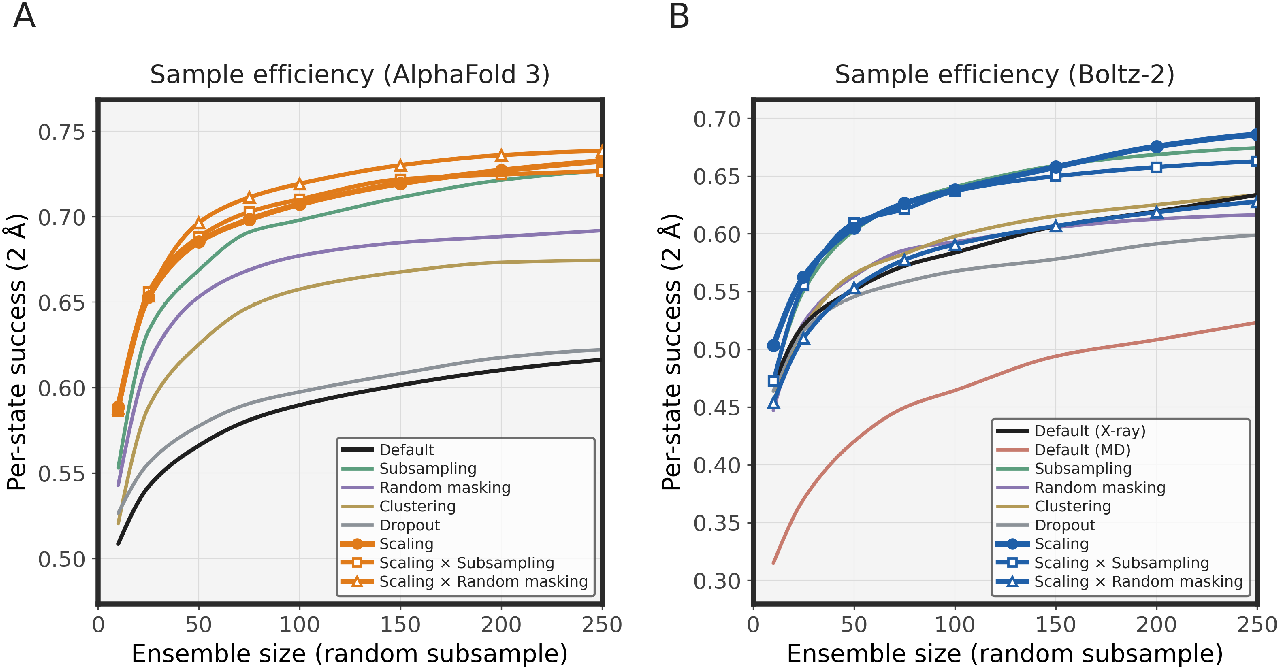
Sample efficiency. Per-state success at 2 Å as a function of ensemble size, drawn at random from the full ensemble, for **(A)** AlphaFold 3 and **(B)** Boltz-2. Default inference, the sampling baselines, scaling, and the two scaling combinations are all compared at equal ensemble sizes.

**Figure S3.**
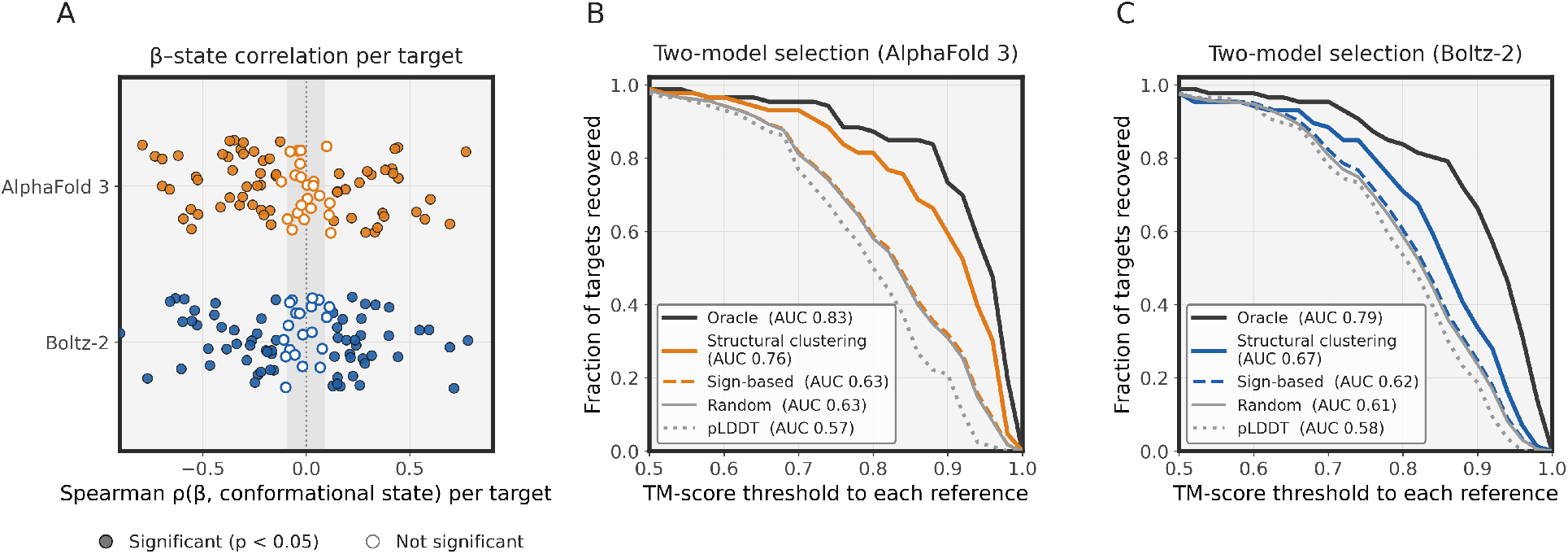
β–state correlation and reference-free model selection. **(A)** Per-target Spearman correlation between *β* and the TM-score difference to the two reference structures. Filled points are significant (*p <* 0.05), and the shaded band marks the non-significant range. **(B, C)** Reference-free selection of two models from the scaled ensemble for AlphaFold 3 (B) and Boltz-2 (C). Success at a TM-score threshold means the two selected models include one within that threshold of each reference. Rules: oracle, structural clustering of secondary-structure-region distance maps, sign-based selection, random baseline, and two highest-pLDDT models.

### S4 Distogram–experiment agreement across targets

Figure S4 shows, for every AlphaFold 3 target with a recovery improvement, the in-range Pearson correlation between the distogram change under scaling and the experimental distance change between the two reference states. The three targets used in Figure 5E–G are marked. The agreement is positive for 34 of the 35 targets (median *r* = 0.56), so the examples are representative of the full set.

**Figure S4.**
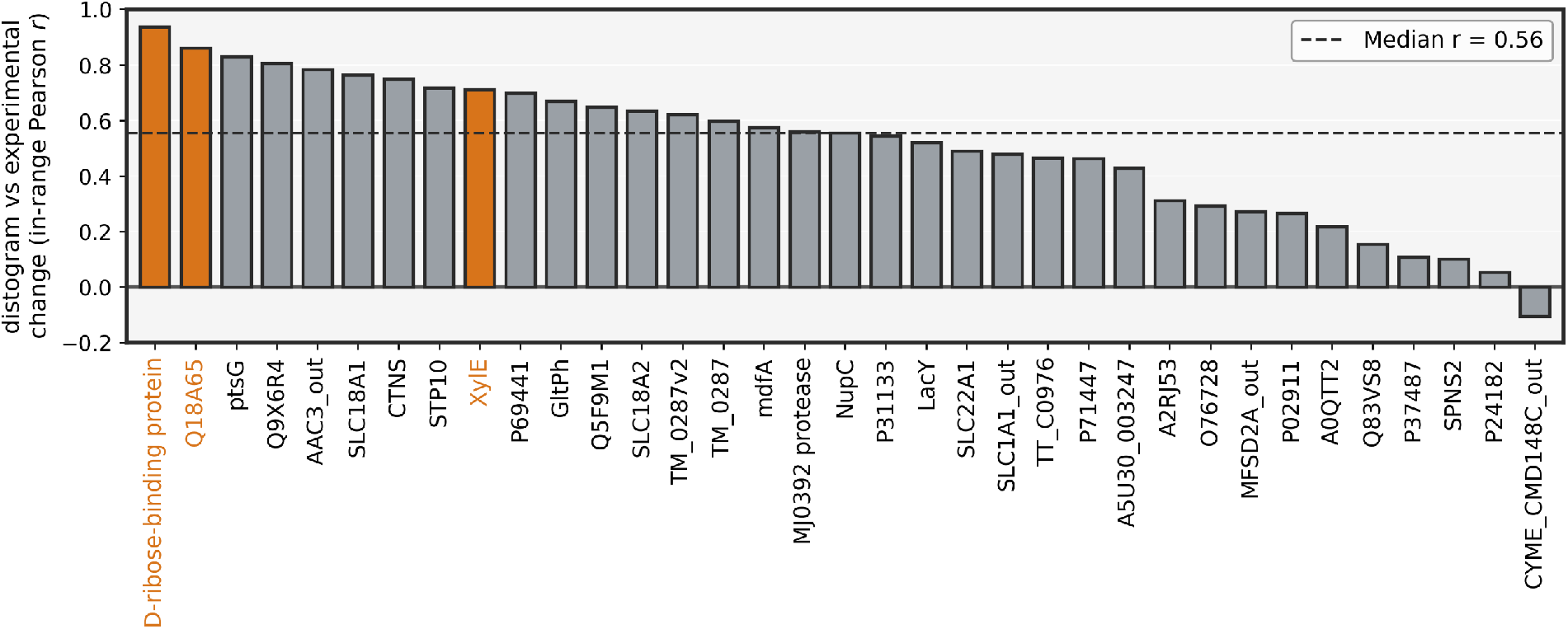
Distogram–experiment agreement for every AlphaFold 3 target with a recovery improvement. For each target, the bar is the in-range Pearson correlation between the per-residue-pair distogram change under scaling (at the target’s *β*^&^) and the experimental distance change between its two reference structures. The three Figure 5 examples are highlighted. The dashed line is the median.

### S5 Per-target supporting figures

#### S5.1 Conformational sampling

For every benchmark target, Figures S5–S10 resolve the conformational sampling of both predictors to individual models. Each target is shown as four panels, from left to right AlphaFold 3 under default inference, AlphaFold 3 under scaling, Boltz-2 under default inference, and Boltz-2 under scaling. Each panel plots the TM-score of every model to the first reference against its TM-score to the second, coloured by *β*, so the spread of an ensemble between the two states is visible directly. Targets are grouped by conformational-change class and the axis range is set per target.

#### S5.2 Distogram bimodality

For each of the 86 benchmark targets, Figures S11–S16 show the AlphaFold 3 distogram at one representative change-contact pair across the 11 values of *β*. For each target the pair was chosen as the change-contact pair maximising the number of *β* values at which the pair is classified as bimodal with fitted modes matching both reference distances, with ties broken by the largest reference distance change. If no *β* value gives a matched bimodal classification the pair with the largest reference distance change is shown. Background colour encodes the classification at each *β* as in Figure 6C, pink for bimodal with peaks matching both reference distances, lavender for bimodal but unmatched, and grey for unimodal.

**Figure S5.**
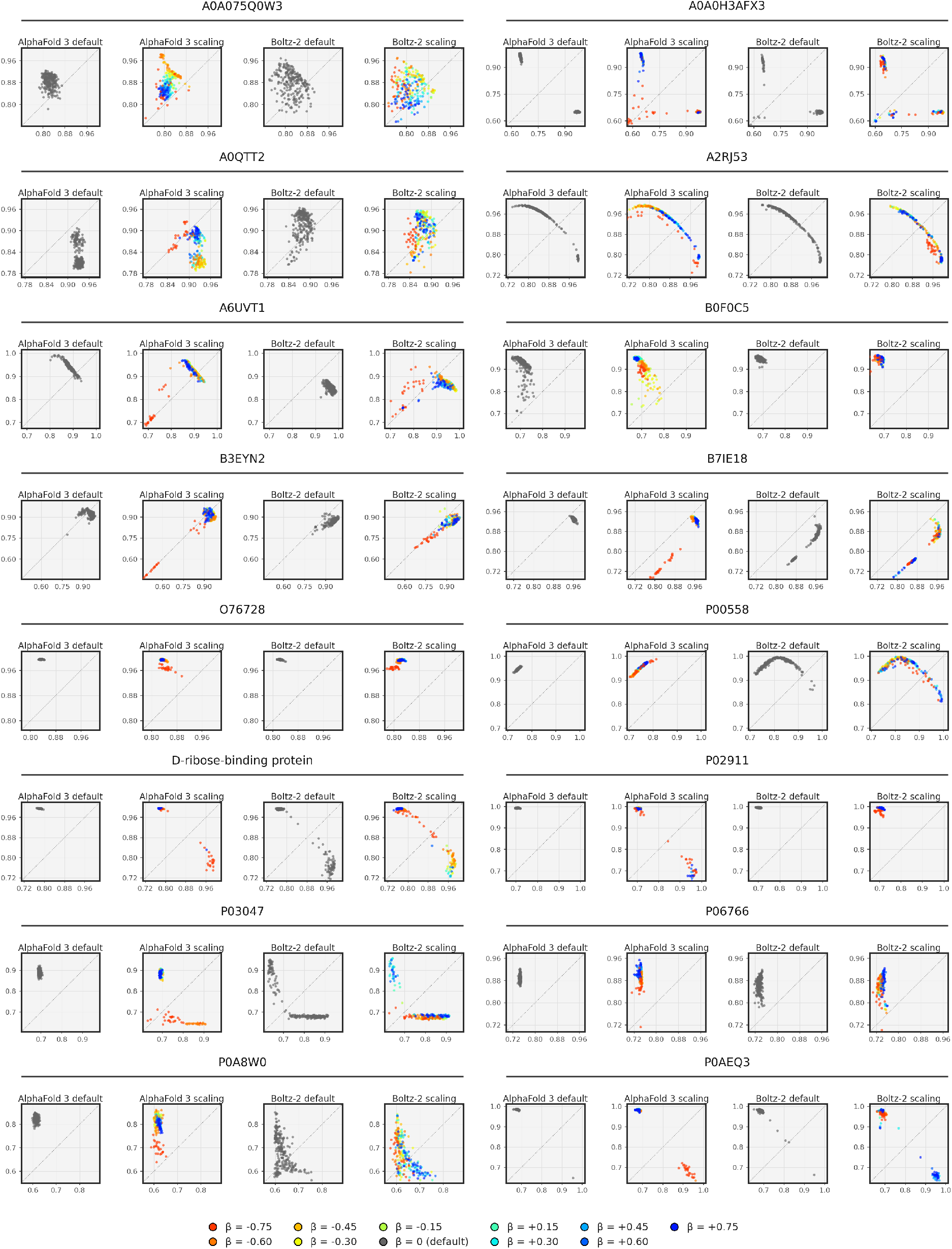
Per-target TM-score scatter, 1 of 6 (domain-motion targets). Each target is shown as four panels, namely AlphaFold 3 and Boltz-2 under default inference and under pair representation scaling, of the TM-score to reference 1 against the TM-score to reference 2, with points coloured by *β* (*β* = 0 is default inference).

**Figure S6.**
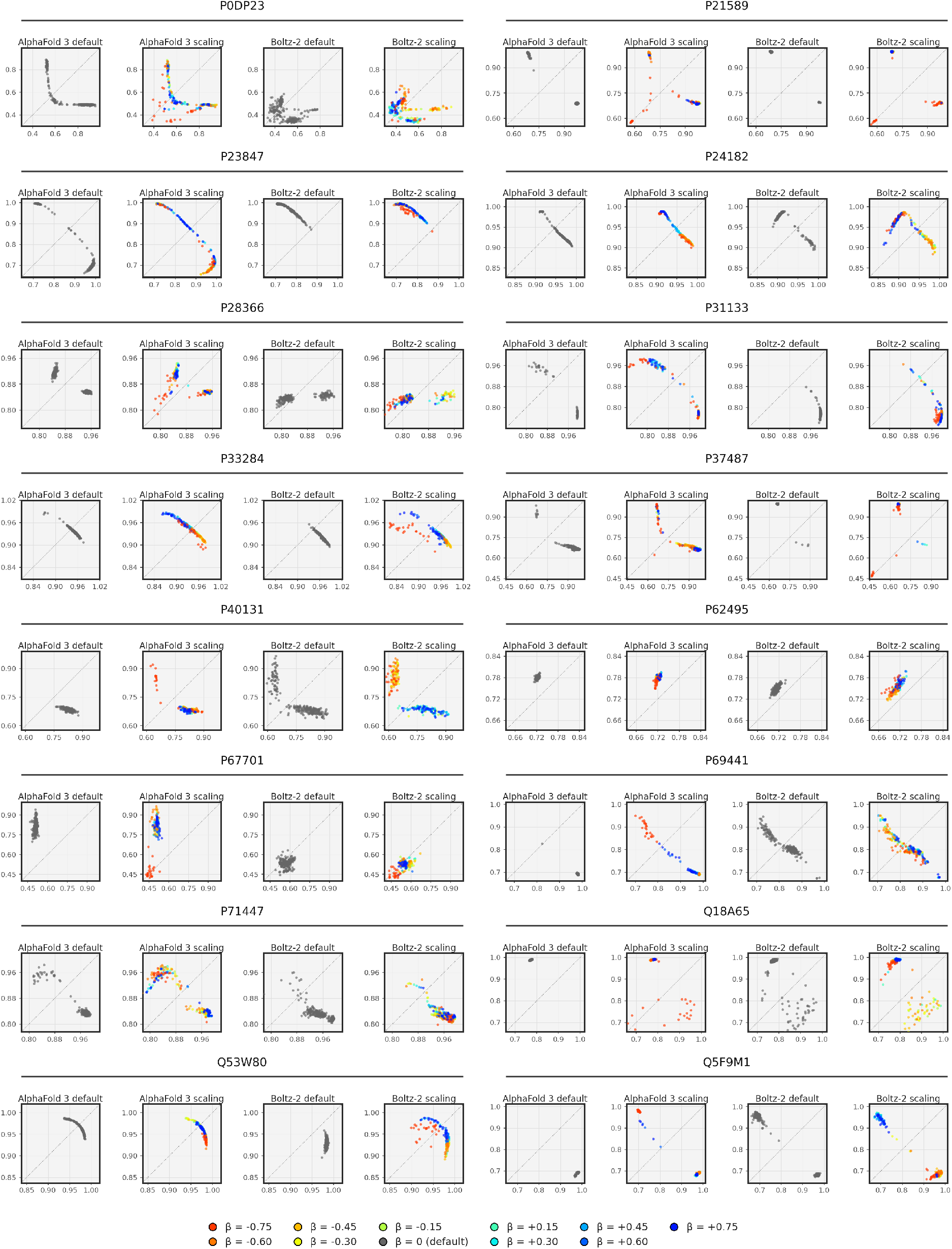
Per-target TM-score scatter, 2 of 6 (domain-motion targets). Panels as in Figure S5.

**Figure S7.**
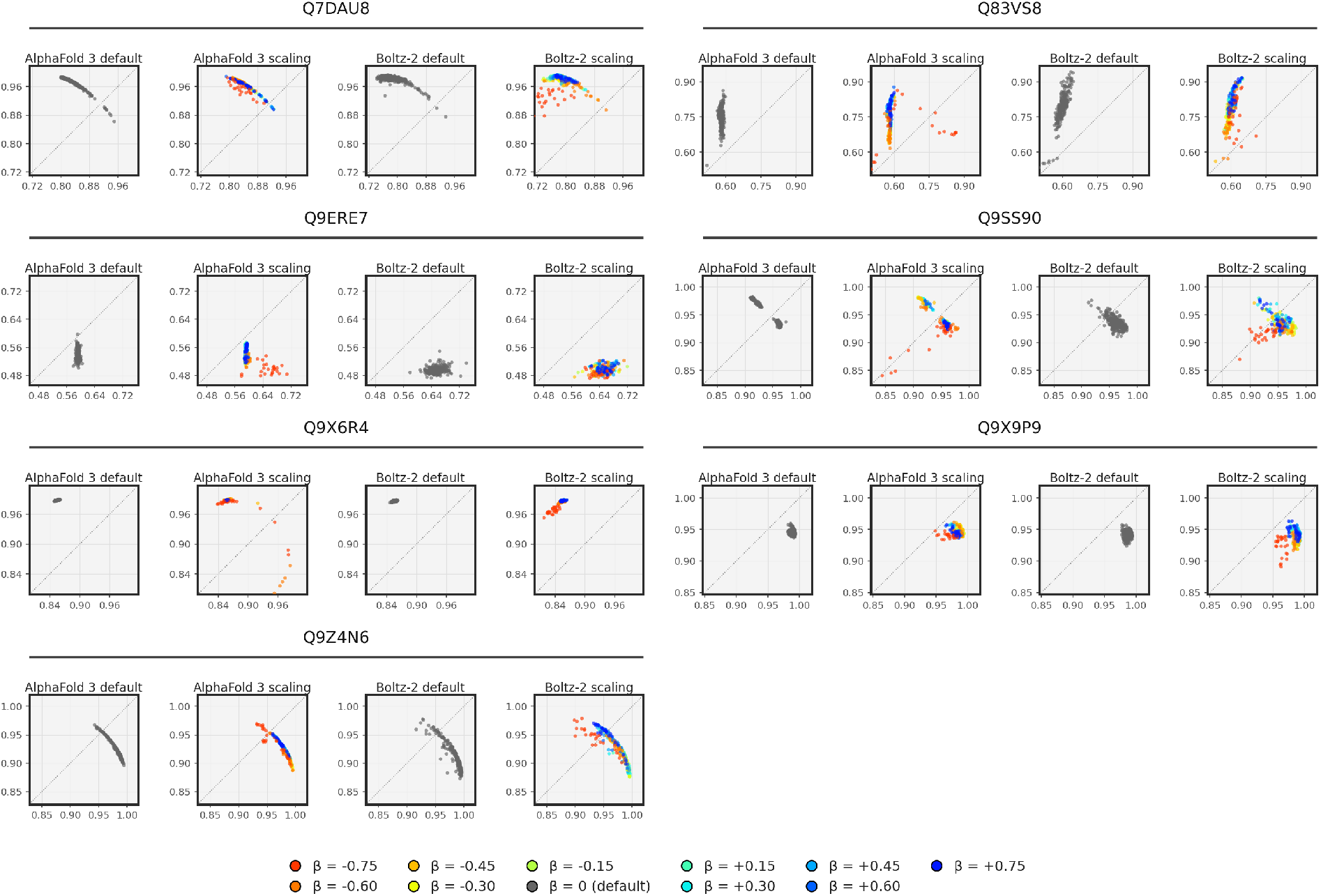
Per-target TM-score scatter, 3 of 6 (domain-motion targets). Panels as in Figure S5.

**Figure S8.**
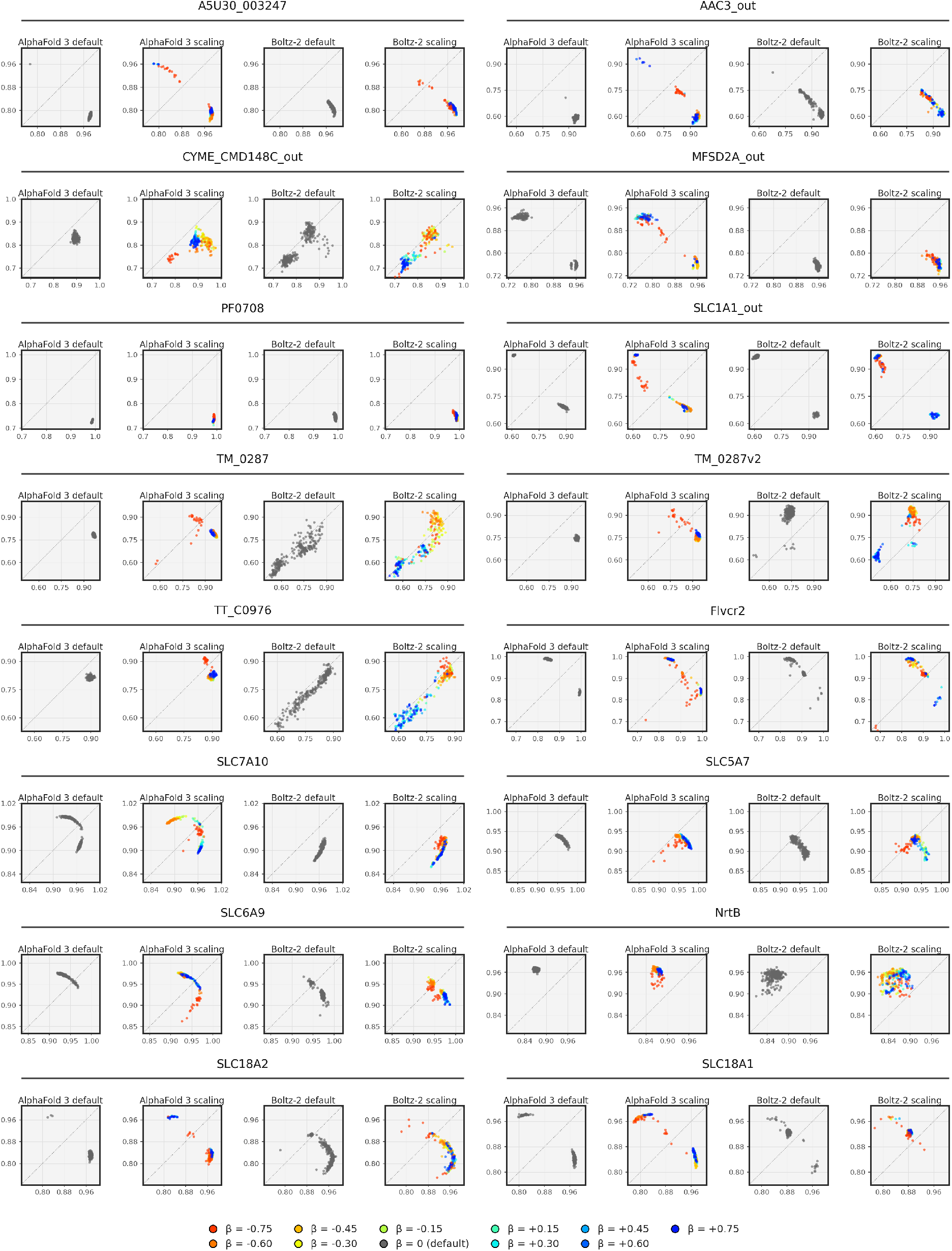
Per-target TM-score scatter, 4 of 6 (transporter targets). Panels as in Figure S5.

**Figure S9.**
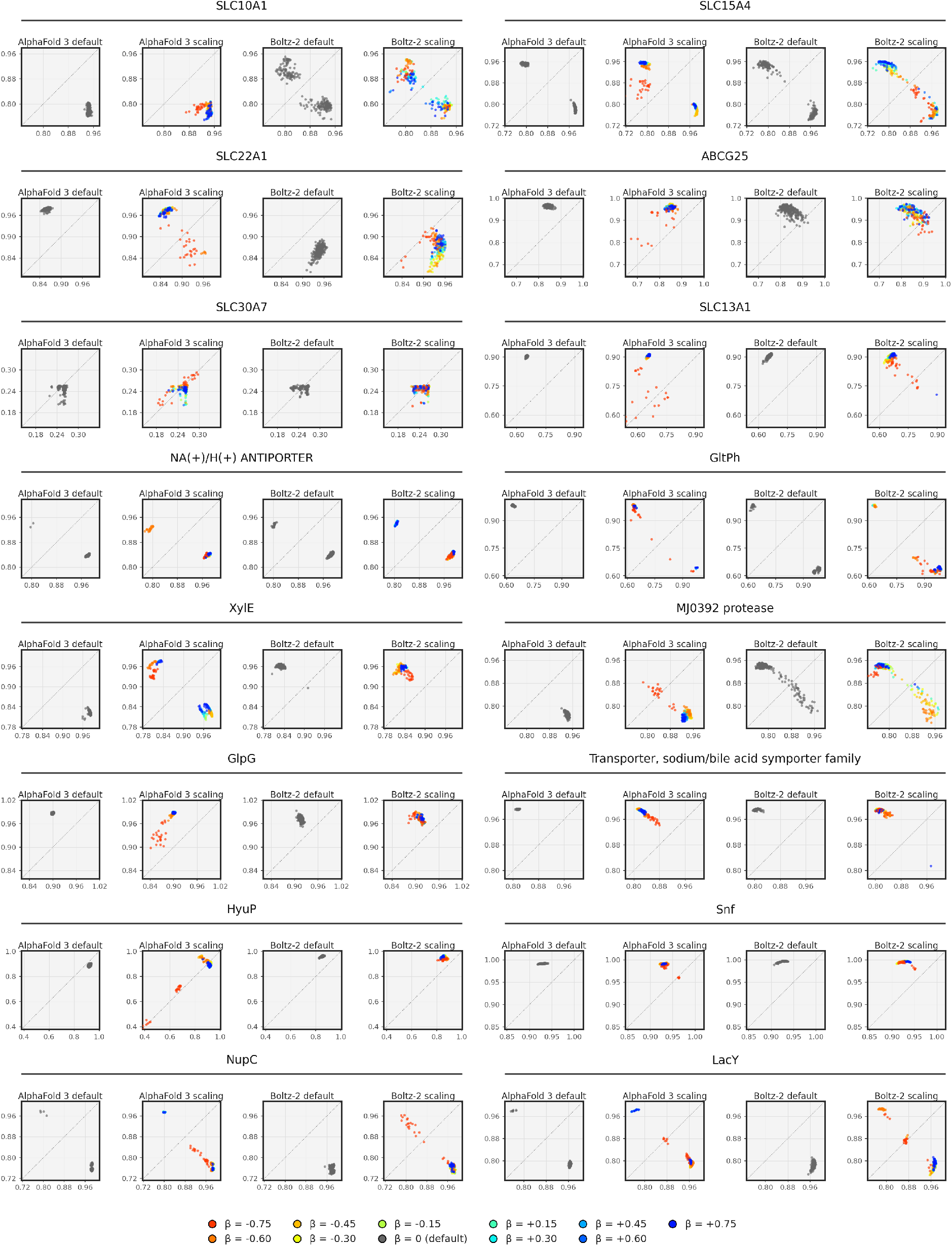
Per-target TM-score scatter, 5 of 6 (transporter targets). Panels as in Figure S5.

**Figure S10.**
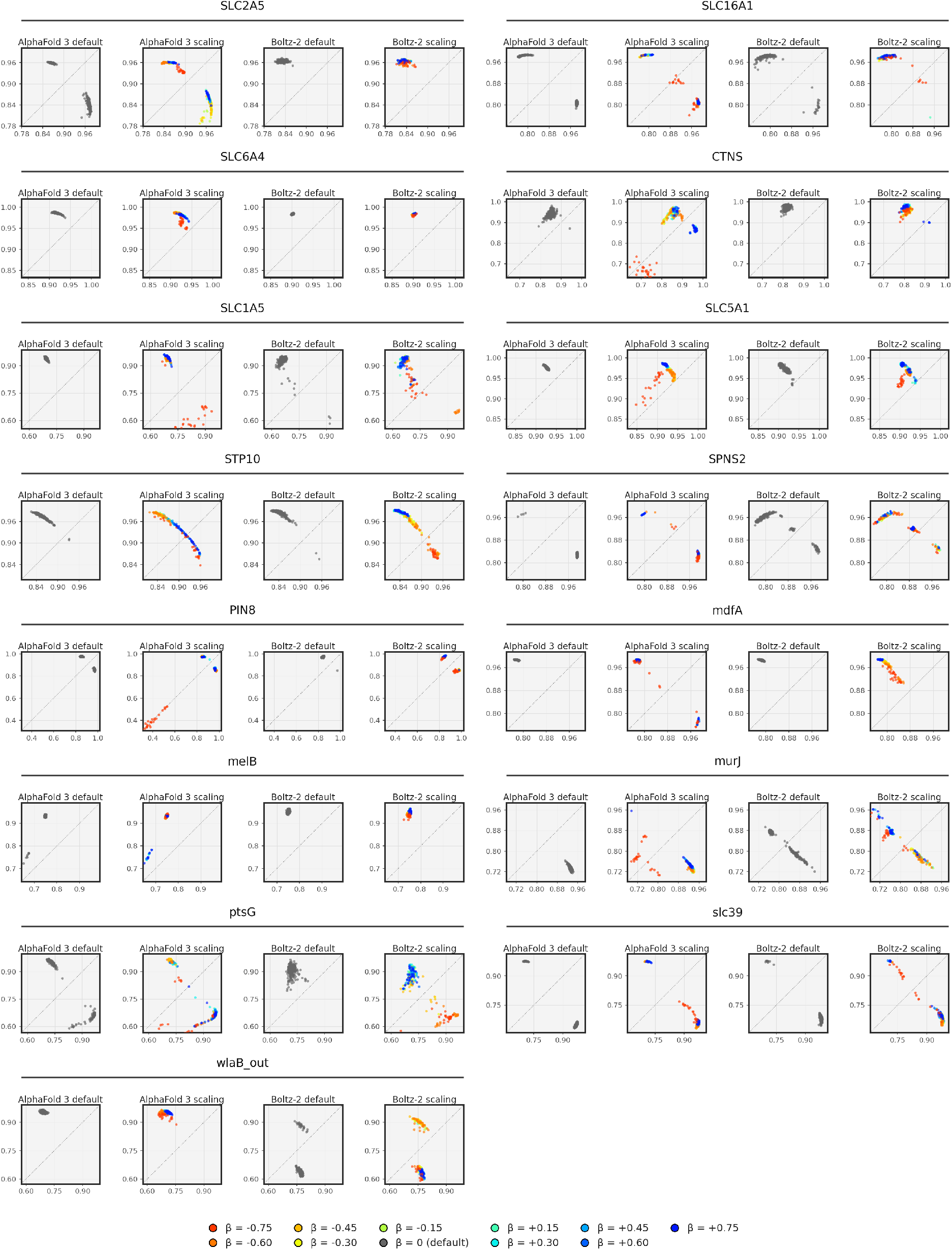
Per-target TM-score scatter, 6 of 6 (transporter targets). Panels as in Figure S5.

**Figure S11.**
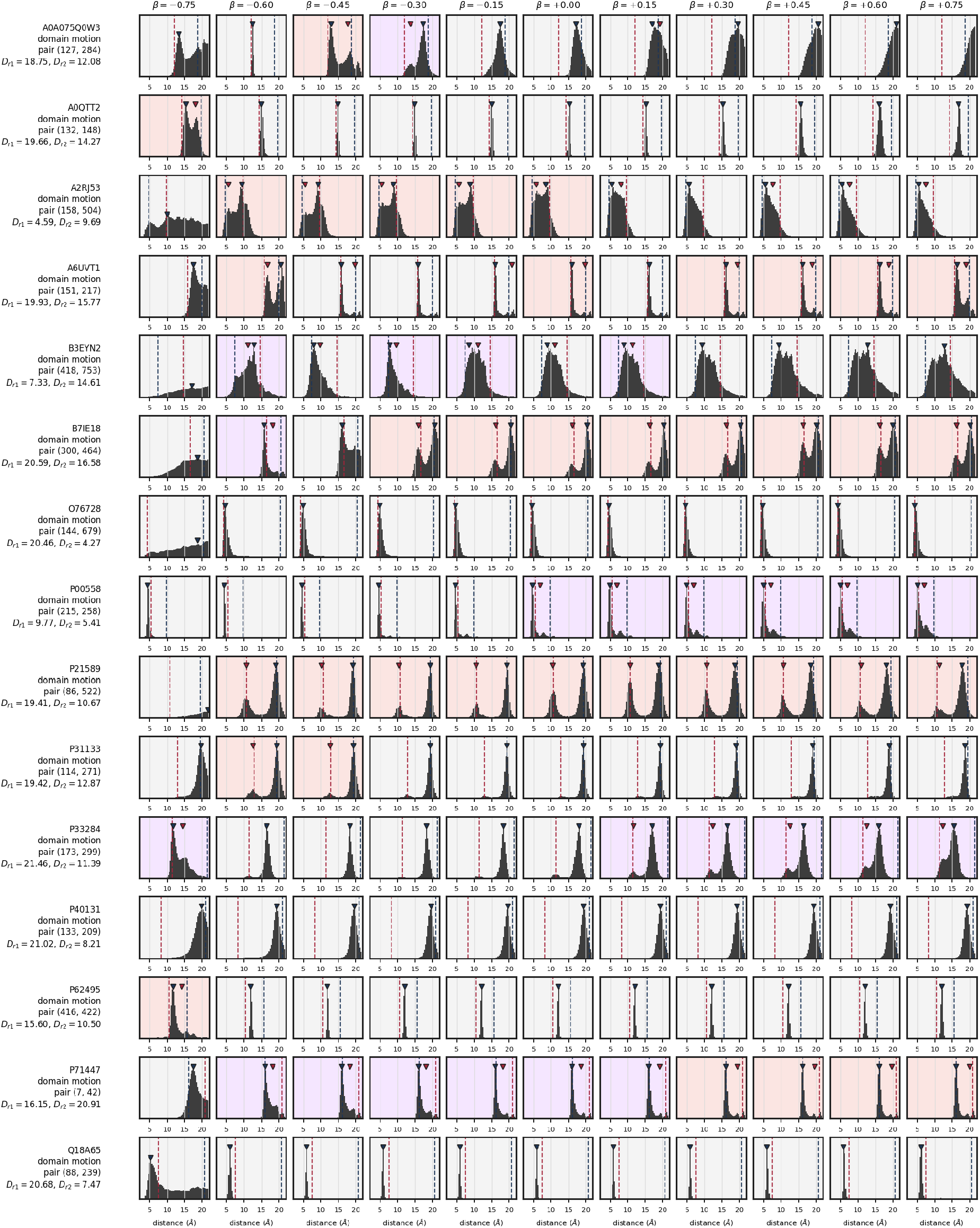
Per-target distogram bimodality, page 1 of 6. Each row is one target and shows the AlphaFold 3 distogram at the representative change-contact pair across the 11 values of *β* (selection criterion in text).

**Figure S12.**
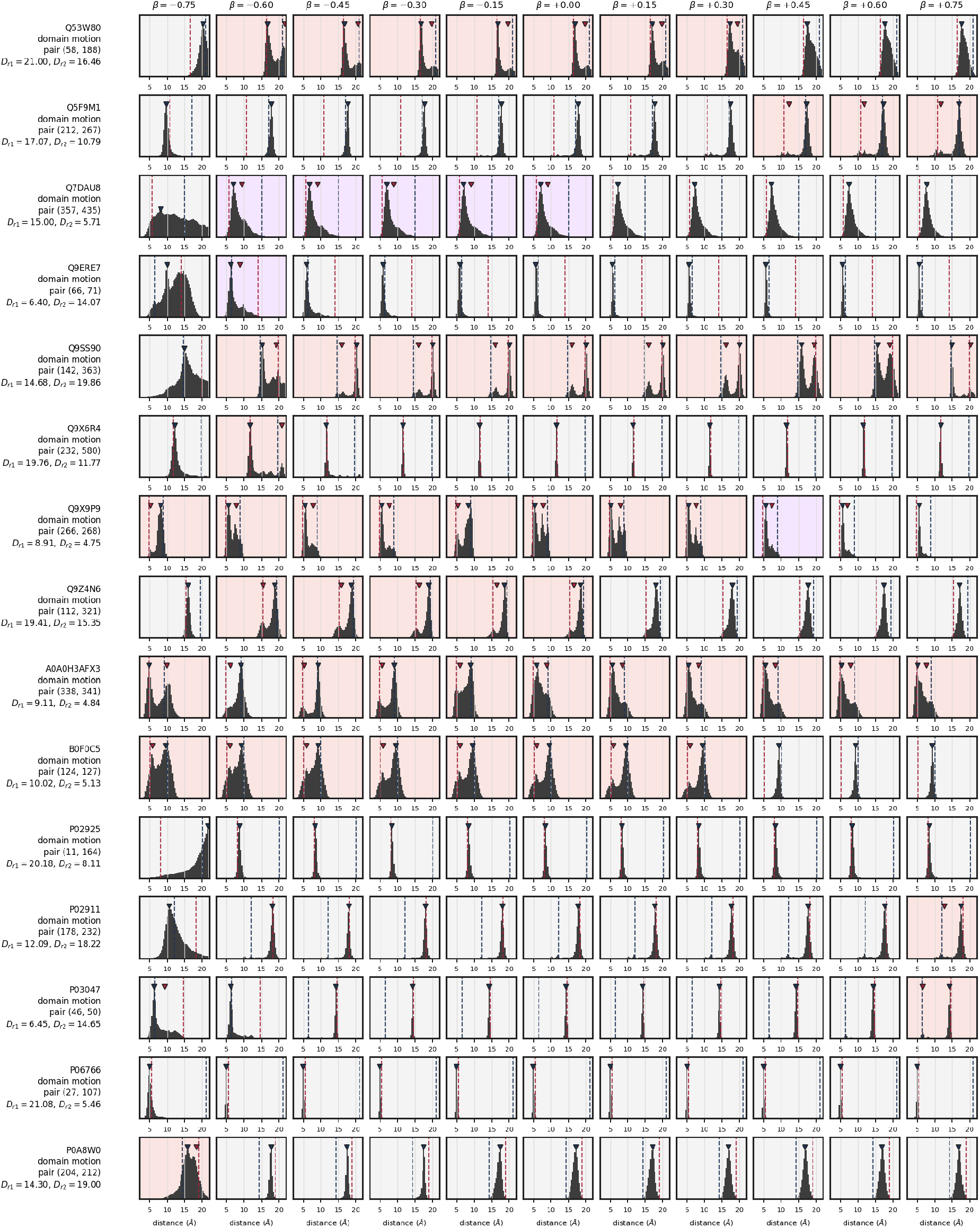
Per-target distogram bimodality, page 2 of 6. Panels as in Figure S11.

**Figure S13.**
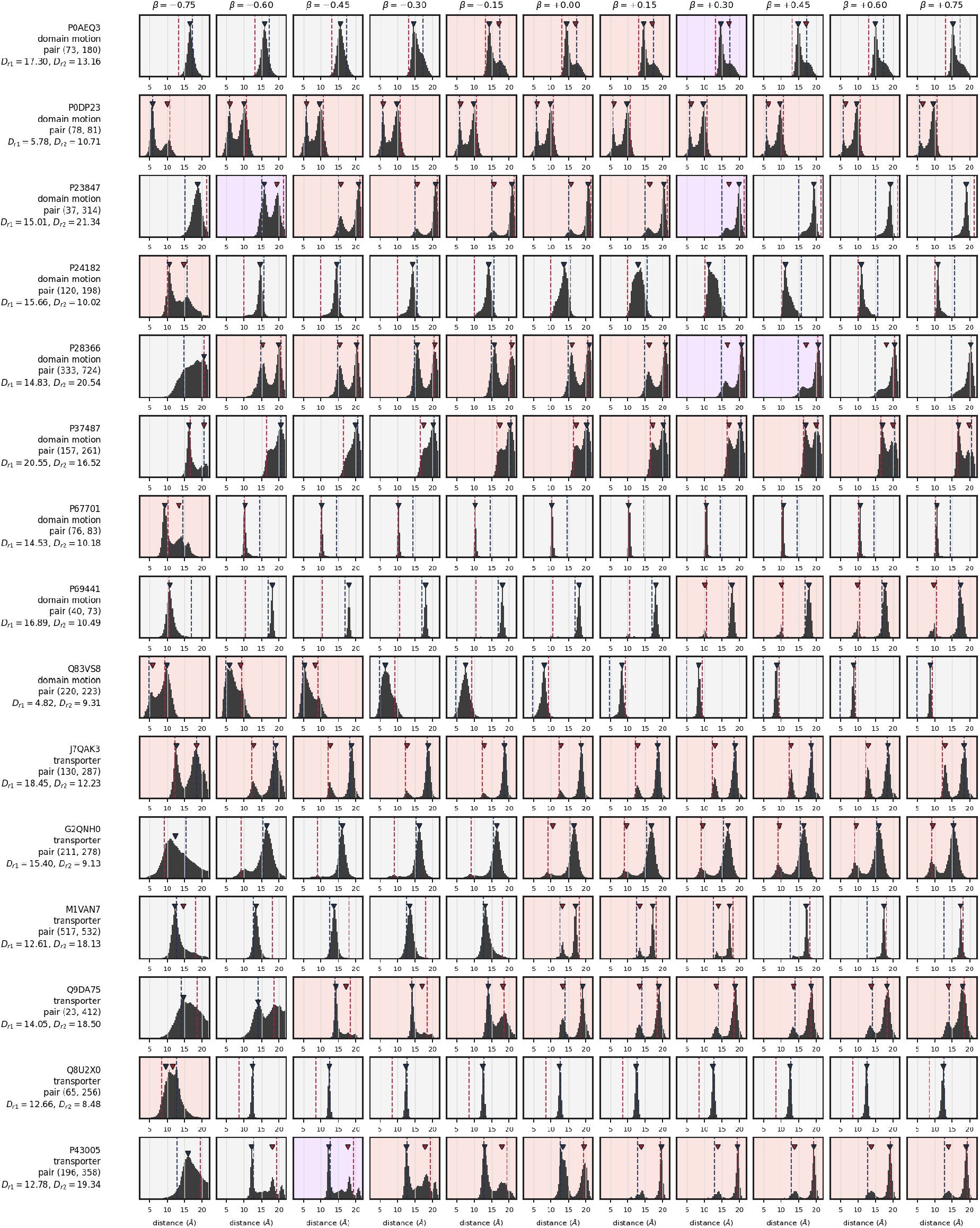
Per-target distogram bimodality, page 3 of 6. Panels as in Figure S11.

**Figure S14.**
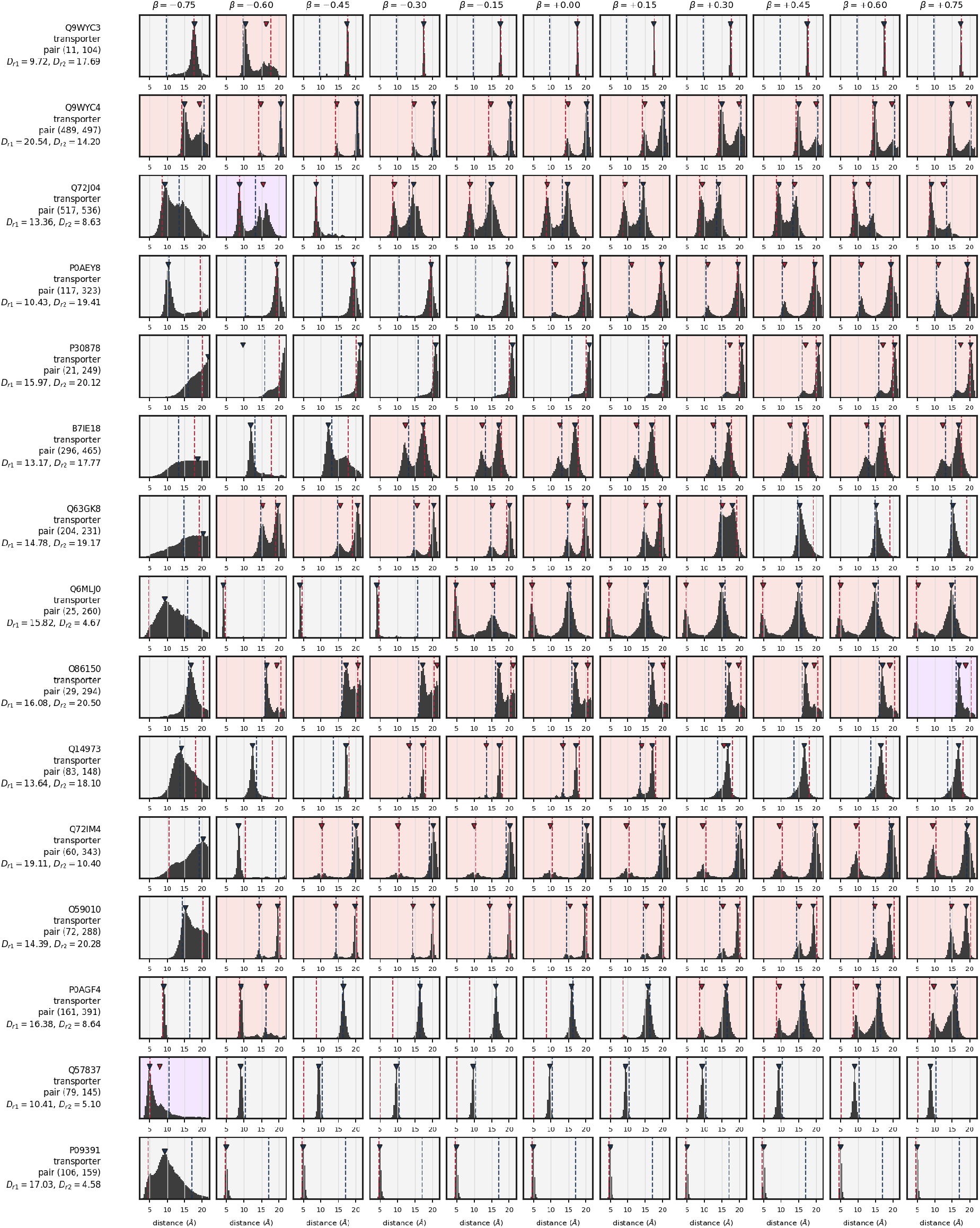
Per-target distogram bimodality, page 4 of 6. Panels as in Figure S11.

**Figure S15.**
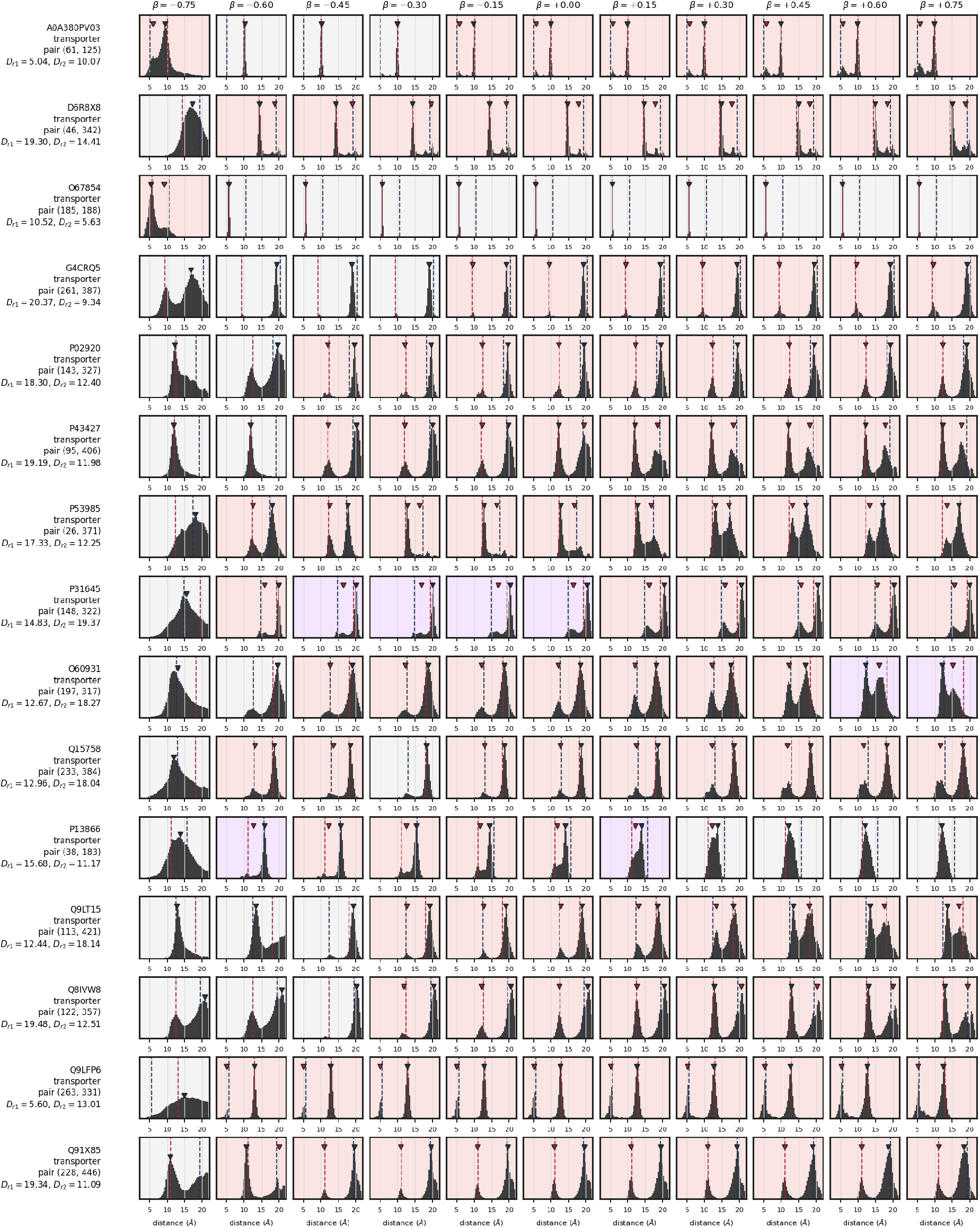
Per-target distogram bimodality, page 5 of 6. Panels as in Figure S11.

**Figure S16.**
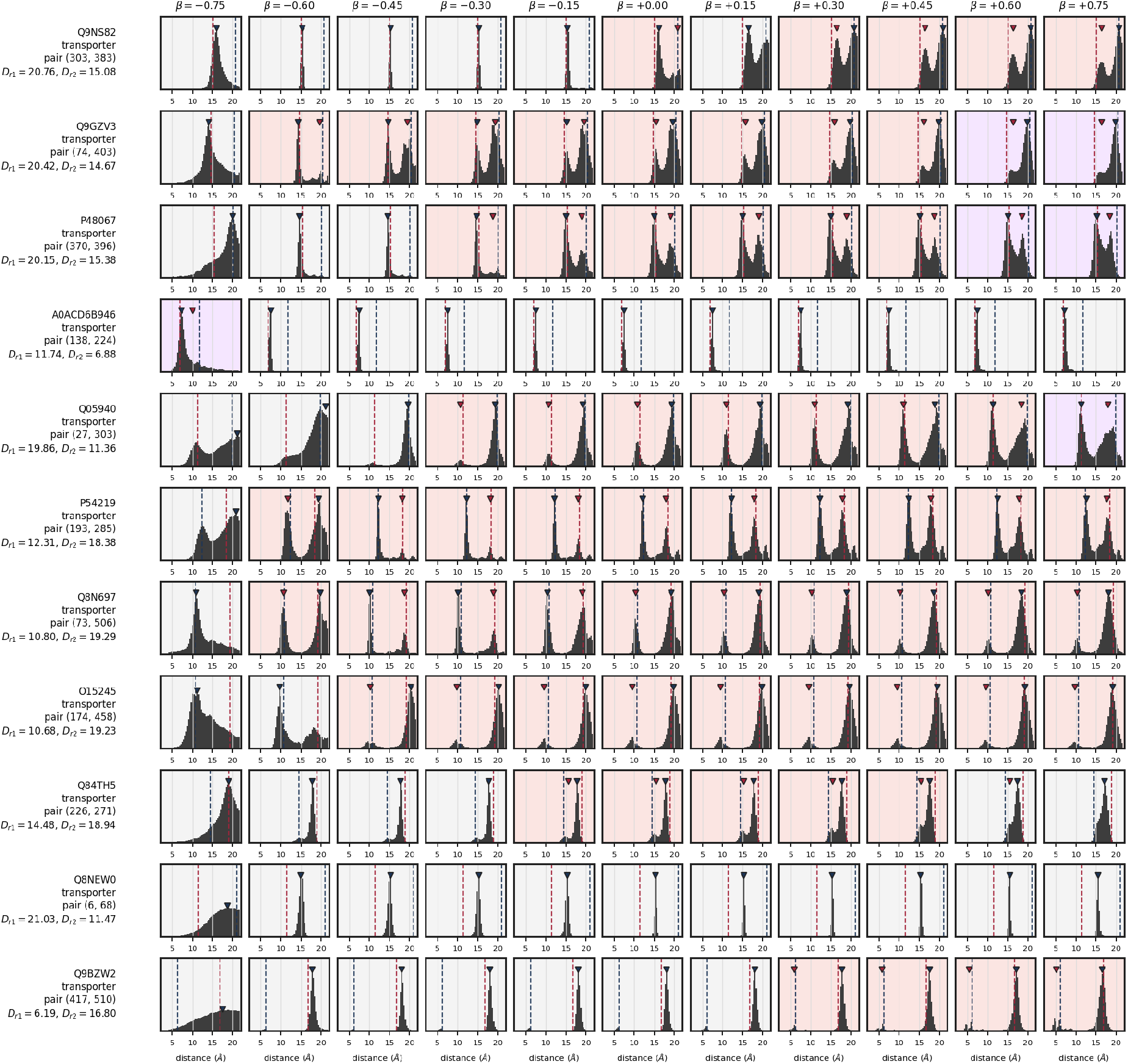
Per-target distogram bimodality, page 6 of 6. Panels as in Figure S11.

## References

[1] John Jumper, Richard Evans, Alexander Pritzel, Tim Green, Michael Figurnov, Olaf Ronneberger, Kathryn Tunyasuvunakool, Russ Bates, Augustin Žídek, Anna Potapenko, Alex Bridgland, Clemens Meyer, Simon A A Kohl, Andrew J Ballard, Andrew Cowie, Bernardino Romera-Paredes, Stanislav Nikolov, Rishub Jain, Jonas Adler, Trevor Back, Stig Petersen, David Reiman, Ellen Clancy, Michal Zielinski, Martin Steinegger, Michalina Pacholska, Tamas Berghammer, Sebastian Bodenstein, David Silver, Oriol Vinyals, Andrew W Senior, Koray Kavukcuoglu, Pushmeet Kohli, and Demis Hassabis. Highly accurate protein structure prediction with AlphaFold. Nature, 596(7873):583–589, August 2021. doi:10.1038/s41586-021-03819-2.

[2] Prajna Mishra and Santosh Kumar Jha. The native state conformational heterogeneity in the energy landscape of protein folding. Biophysical Chemistry, 283:106761, 2022. ISSN 0301-4622. 10.1016/j.bpc.2022.106761. URL https://www.sciencedirect.com/science/article/pii/S0301462222000035.

[3] Gregory R. Bowman. Alphafold and protein folding: Not dead yet! the frontier is conformational ensembles. Annual Review of Biomedical Data Science, 7(1):51–57, 2024. doi:10.1146/annurev-biodatasci-102423-011435.

[4] Xinyue Cui, Lingyu Ge, Xia Chen, Zexin Lv, Suhui Wang, Xiaogen Zhou, and Guijun Zhang. Beyond static structures: protein dynamic conformations modeling in the post-AlphaFold era. Briefings in Bioinformatics, 26(4): bbaf340, 2025. doi:10.1093/bib/bbaf340.

[5] Richard A Stein and Hassane S Mchaourab. SPEACH_AF: Sampling protein ensembles and conformational hetero-geneity with Alphafold2. PLoS Comput. Biol., 18(8):e1010483, August 2022. doi:10.1371/journal.pcbi.1010483.

[6] Diego Del Alamo, Davide Sala, Hassane S Mchaourab, and Jens Meiler. Sampling alternative conformational states of transporters and receptors with AlphaFold2. Elife, 11, March 2022. doi:10.7554/eLife.75751.

[7] Gabriel Monteiro da Silva, Jennifer Y. Cui, David C. Dalgarno, George P. Lisi, and Brenda M. Rubenstein. High-throughput prediction of protein conformational distributions with subsampled alphafold2. Nature Communications, 15(1):2464, 2024. doi:10.1038/s41467-024-46715-9.

[8] Elaine Tao and Ben Corry. AlphaFold2 captures conformational transitions in the voltage-gated channel superfamily. Biophysical Journal, 124(19):3291–3303, March 2025. doi:10.1016/j.bpj.2025.08.033.

[9] Yogesh Kalakoti and Björn Wallner. AFsample2 predicts multiple conformations and ensembles with AlphaFold2. *Commun*. Biol., 8(1):373, March 2025. doi:10.1038/s42003-025-07791-9.

[10] Yogesh Kalakoti and Björn Wallner. AFsample3: Generating and selecting multiple conformational states with AlphaFold3. bioRxiv, 2026. doi:10.64898/2026.01.16.699904.

[11] Hannah K Wayment-Steele, Adedolapo Ojoawo, Renee Otten, Julia M Apitz, Warintra Pitsawong, Marc Hömberger, Sergey Ovchinnikov, Lucy Colwell, and Dorothee Kern. Predicting multiple conformations via sequence clustering and AlphaFold2. Nature, 625(7996):832–839, January 2024. doi:10.1038/s41586-023-06832-9.

[12] Björn Wallner. AFsample: improving multimer prediction with AlphaFold using massive sampling. Bioinformatics, 39(9):btad573, September 2023. doi:10.1093/bioinformatics/btad573.

[13] David Wu and Liang Feng. Robust prediction of multiple protein conformations with entropy guidance. bioRxiv, page 2025.04.26.650728, April 2025. doi:10.1101/2025.04.26.650728.

[14] Francesca Peccati, Sara Alunno-Rufini, and Gonzalo Jiménez-Osés. Accurate prediction of enzyme thermostabilization with rosetta using alphafold ensembles. Journal of Chemical Information and Modeling, 63(3):898–909, 2023. doi:10.1021/acs.jcim.2c01083.

[15] Bodhi P. Vani, Akashnathan Aranganathan, and Pratyush Tiwary. Exploring kinase asp-phe-gly (dfg) loop conformational stability with alphafold2-rave. Journal of Chemical Information and Modeling, 64(7):2789–2797, 2024. doi:10.1021/acs.jcim.3c01436.

[16] Ruth Nussinov, Mingzhen Zhang, Yonglan Liu, and Hyunbum Jang. AlphaFold, artificial intelligence (AI), and allostery. J. Phys. Chem. B, 126(34):6372–6383, September 2022. doi:10.1021/acs.jpcb.2c04346.

[17] Josh Abramson, Jonas Adler, Jack Dunger, Richard Evans, Tim Green, Alexander Pritzel, Olaf Ronneberger, Lindsay Willmore, Andrew J Ballard, Joshua Bambrick, Sebastian W Bodenstein, David A Evans, Chia-Chun Hung, Michael O’Neill, David Reiman, Kathryn Tunyasuvunakool, Zachary Wu, Akvilė Žemgulytė, Eirini Arvaniti, Charles Beattie, Ottavia Bertolli, Alex Bridgland, Alexey Cherepanov, Miles Congreve, Alexander I Cowen-Rivers, Andrew Cowie, Michael Figurnov, Fabian B Fuchs, Hannah Gladman, Rishub Jain, Yousuf A Khan, Caroline M R Low, Kuba Perlin, Anna Potapenko, Pascal Savy, Sukhdeep Singh, Adrian Stecula, Ashok Thillaisundaram, Catherine Tong, Sergei Yakneen, Ellen D Zhong, Michal Zielinski, Augustin Žídek, Victor Bapst, Pushmeet Kohli, Max Jaderberg, Demis Hassabis, and John M Jumper. Accurate structure prediction of biomolecular interactions with AlphaFold 3. Nature, 630(8016):493–500, June 2024. doi:10.1038/s41586-024-07487-w.

[18] Jeremy Wohlwend, Gabriele Corso, Saro Passaro, Noah Getz, Mateo Reveiz, Ken Leidal, Wojtek Swiderski, Liam Atkinson, Tally Portnoi, Itamar Chinn, Jacob Silterra, Tommi Jaakkola, and Regina Barzilay. Boltz-1 democratizing biomolecular interaction modeling. bioRxiv, November 2024. doi:10.1101/2024.11.19.624167.

[19] Saro Passaro, Gabriele Corso, Jeremy Wohlwend, Mateo Reveiz, Stephan Thaler, Vignesh Ram Somnath, Noah Getz, Tally Portnoi, Julien Roy, Hannes Stark, David Kwabi-Addo, Dominique Beaini, Tommi Jaakkola, and Regina Barzilay. Boltz-2: Towards accurate and efficient binding affinity prediction. bioRxiv, page 2025.06.14.659707, June 2025. doi:10.1101/2025.06.14.659707.

[20] Jonathan Ho, Ajay Jain, and Pieter Abbeel. Denoising diffusion probabilistic models, 2020. URL https://arxiv.org/abs/2006.11239.

[21] G V T Swapna, N Dube, M J Roth, and G T Montelione. Memorization bias impacts modeling of alternative conformational states of solute carrier membrane proteins with methods from deep learning. PLOS Computational Biology, 21(10):e1013590, 2025. doi:10.1371/journal.pcbi.1013590.

[22] Devlina Chakravarty, Myeongsang Lee, and Lauren L Porter. Proteins with alternative folds reveal blind spots in AlphaFold-based protein structure prediction. Curr. Opin. Struct. Biol., 90(102973):102973, February 2025. doi:10.1016/j.sbi.2024.102973.

[23] Sarah Lewis, Tim Hempel, José Jiménez-Luna, Michael Gastegger, Yu Xie, Andrew Y. K. Foong, Victor García Satorras, Osama Abdin, Bastiaan S. Veeling, Iryna Zaporozhets, Yaoyi Chen, Soojung Yang, Adam E. Foster, Arne Schneuing, Jigyasa Nigam, Federico Barbero, Vincent Stimper, Andrew Campbell, Jason Yim, Marten Lienen, Yu Shi, Shuxin Zheng, Hannes Schulz, Usman Munir, Roberto Sordillo, Ryota Tomioka, Cecilia Clementi, and Frank Noé. Scalable emulation of protein equilibrium ensembles with generative deep learning. Science, 389 (6761):eadv9817, 2025. doi:10.1126/science.adv9817.

[24] Till Siebenmorgen, Filipe Menezes, Sabrina Benassou, Erinc Merdivan, Kieran Didi, André Santos Dias Mourão, Radosław Kitel, Pietro Liò, Stefan Kesselheim, Marie Piraud, Fabian J. Theis, Michael Sattler, and Grzegorz M. Popowicz. MISATO: machine learning dataset of protein–ligand complexes for structure-based drug discovery. Nature Computational Science, 4(5):367–378, 2024. doi:10.1038/s43588-024-00627-2.

[25] Yann Vander Meersche, Gabriel Cretin, Aria Gheeraert, Jean-Christophe Gelly, and Tatiana Galochkina. ATLAS: protein flexibility description from atomistic molecular dynamics simulations. Nucleic Acids Research, 52(D1): D384–D392, 2024. doi:10.1093/nar/gkad1084.

[26] Antonio Mirarchi, Toni Giorgino, and Gianni De Fabritiis. mdCATH: A large-scale MD dataset for data-driven computational biophysics. Scientific Data, 11(1):1299, 2024. doi:10.1038/s41597-024-04140-z.

[27] Jiaxuan Li, Zefeng Zhu, and Chen Song. Predicting the alternative conformation of a known protein structure based on the distance map of alphafold2. *bioRxiv*, 2024. doi:10.1101/2024.06.09.598121.

[28] Büşra Savaş, Ayşe Berçin Barlas, and Ezgi Karaca. Exploring the potential of AlphaFold distograms for predicting binding-induced hinge motions. FEBS Letters, 600(10):1558–1570, 2026. doi:10.1002/1873-3468.70297.

[29] Jonathan Feldman and Jeffrey Skolnick. Alphainterp: Mechanistic interpretability of alphafold 3 reveals how evolutionary information shapes protein structure prediction. *bioRxiv*, 2026. doi:10.64898/2026.04.22.720175.

[30] Zeming Lin, Halil Akin, Roshan Rao, Brian Hie, Zhongkai Zhu, Wenting Lu, Nikita Smetanin, Robert Verkuil, Ori Kabeli, Yaniv Shmueli, Allan dos Santos Costa, Maryam Fazel-Zarandi, Tom Sercu, Salvatore Candido, and Alexander Rives. Evolutionary-scale prediction of atomic-level protein structure with a language model. Science, 379(6637):1123–1130, 2023. doi:10.1126/science.ade2574.

[31] Kevin Lu, Jannik Brinkmann, Stefan Huber, Aaron Mueller, Yonatan Belinkov, David Bau, and Chris Wendler. Two stages of folding: Convergent mechanisms in AI protein folding trunks. *arXiv preprint arXiv:2602.06020*, 2026. doi:10.48550/arXiv.2602.06020.

[32] Myeongsang Lee, Joseph W Schafer, Jeshuwin Prabakaran, Devlina Chakravarty, Madeleine F Clore, and Lauren L Porter. Large-scale predictions of alternative protein conformations by AlphaFold2-based sequence association. Nat. Commun., 16(1):5622, July 2025. doi:10.1038/s41467-025-60759-5.

[33] Gonzalo Jiménez-Osés and Francesca Peccati. Structure prediction of alternate frame folding systems with alphafold3. Journal of Chemical Information and Modeling, 65(15):8229–8237, August 2025. doi:10.1021/acs.jcim.5c00906. URL https://pubmed.ncbi.nlm.nih.gov/40785361/.

[34] Björn Wallner, Alexey Amunts, Andreas Naschberger, Björn Nystedt, and Claudio Mirabello. dgram2dmap: Extraction, visualisation and formatting of distance constraints from alphafold distograms. *bioRxiv*, 2022. doi:10.1101/2022.12.08.519560.

[35] Yan Wang, Lihao Wang, Yuning Shen, Yiqun Wang, Huizhuo Yuan, Yue Wu, and Quanquan Gu. Protein conformation generation via force-guided SE(3) diffusion models. In Ruslan Salakhutdinov, Zico Kolter, Katherine Heller, Adrian Weller, Nuria Oliver, Jonathan Scarlett, and Felix Berkenkamp, editors, Proceedings of the 41st International Conference on Machine Learning, volume 235 of *Proceedings of Machine Learning Research*, pages 56835–56859. PMLR, 21–27 Jul 2024. URL https://proceedings.mlr.press/v235/wang24cv.html.

[36] Ameya Daigavane, Bodhi P. Vani, Darcy Davidson, Saeed Saremi, Joshua A. Rackers, and Joseph Kleinhenz. Jamun: Bridging smoothed molecular dynamics and score-based learning for conformational ensembles. *arXiv preprint arXiv:2410.14621*, 2024. doi:10.48550/arXiv.2410.14621. URL https://arxiv.org/abs/2410.14621.

[37] Gustav Olanders, Giulia Testa, Alessandro Tibo, Eva Nittinger, and Christian Tyrchan. Challenge for deep learning: Protein structure prediction of ligand-induced conformational changes at allosteric and orthosteric sites. Journal of Chemical Information and Modeling, 64(22):8481–8494, 2024. doi:10.1021/acs.jcim.4c01475.

[38] Benjamin P Brown, Richard A Stein, Jens Meiler, and Hassane S Mchaourab. Approximating projections of conformational boltzmann distributions with AlphaFold2 predictions: Opportunities and limitations. J. Chem. Theory Comput., 20(3):1434–1447, February 2024. doi:10.1021/acs.jctc.3c01081.

[39] Alex Bateman, Maria-Jesus Martin, Sandra Orchard, Michele Magrane, Shadab Ahmad, Emanuele Alpi, Emily H. Bowler-Barnett, Ramona Britto, Hema Bye-A-Jee, Austra Cukura, Paul Denny, Tunca Dogan, ThankGod Ebenezer, Jun Fan, Penelope Garmiri, Leonardo Jose da Costa Gonzales, Emma Hatton-Ellis, Abdulrahman Hussein, Alexandr Ignatchenko, Giuseppe Insana, Rizwan Ishtiaq, Vishal Joshi, Dushyanth Jyothi, Swaathi Kandasaamy, Antonia Lock, Aurelien Luciani, Marija Lugaric, Jie Luo, Yvonne Lussi, Alistair MacDougall, Fabio Madeira, Mahdi Mahmoudy, Alok Mishra, Katie Moulang, Andrew Nightingale, Sangya Pundir, Guoying Qi, Shriya Raj, Pedro Raposo, Daniel L. Rice, Rabie Saidi, Rafael Santos, Elena Speretta, James Stephenson, Prabhat Totoo, Edward Turner, Nidhi Tyagi, Preethi Vasudev, Kate Warner, Xavier Watkins, Rossana Zaru, Hermann Zellner, Alan J. Bridge, Lucila Aimo, Ghislaine Argoud-Puy, Andrea H. Auchincloss, Kristian B. Axelsen, Parit Bansal, Delphine Baratin, Teresa M. Batista Neto, Marie-Claude Blatter, Jerven T. Bolleman, Emmanuel Boutet, Lionel Breuza, Blanca Cabrera Gil, Cristina Casals-Casas, Kamal Chikh Echioukh, Elisabeth Coudert, Beatrice Cuche, Edouard de Castro, Anne Estreicher, Maria L. Famiglietti, Marc Feuermann, Elisabeth Gasteiger, Pascale Gaudet, Sebastien Gehant, Vivienne Gerritsen, Arnaud Gos, Nadine Gruaz, Chantal Hulo, Nevila Hyka-Nouspikel, Florence Jungo, Arnaud Kerhornou, Philippe Le Mercier, Damien Lieberherr, Patrick Masson, Anne Morgat, Venkatesh Muthukrishnan, Salvo Paesano, Ivo Pedruzzi, Sandrine Pilbout, Lucille Pourcel, Sylvain Poux, Monica Pozzato, Manuela Pruess, Nicole Redaschi, Catherine Rivoire, Christian J. A. Sigrist, Karin Sonesson, Shyamala Sundaram, Cathy H. Wu, Cecilia N. Arighi, Leslie Arminski, Chuming Chen, Yongxing Chen, Hongzhan Huang, Kati Laiho, Peter McGarvey, Darren A. Natale, Karen Ross, C. R. Vinayaka, Qinghua Wang, Yuqi Wang, and Jian Zhang. Uniprot: the universal protein knowledgebase in 2023. Nucleic Acids Research, 51(D1):D523–D531, November 2023. ISSN 1362-4962. doi:10.1093/nar/gkac1052. URL 10.1093/nar/gkac1052.

[40] Helen M. Berman, John Westbrook, Zukang Feng, Gary Gilliland, Talapady N. Bhat, Helge Weissig, Ilya N. Shindyalov, and Philip E. Bourne. The protein data bank. Nucleic Acids Research, 28(1):235–242, 2000. doi:10.1093/nar/28.1.235.

[41] Yogesh Kalakoti and Björn Wallner. Datasets: AFsample2 predicts multiple conformations and ensembles with AlphaFold2, dec 2024. URL 10.5281/zenodo.14534088.

[42] Tengyu Xie and Jing Huang. Can protein structure prediction methods capture alternative conformations of membrane transporters? Journal of Chemical Information and Modeling, 64(8):3524–3536, 2024. doi:10.1021/acs.jcim.3c01936.

[43] Martin Ester, Hans-Peter Kriegel, Jörg Sander, and Xiaowei Xu. A density-based algorithm for discovering clusters in large spatial databases with noise. In Proceedings of the Second International Conference on Knowledge Discovery and Data Mining, KDD’96, page 226–231. AAAI Press, 1996.

[44] Chengxin Zhang, Morgan Shine, Anna Marie Pyle, and Yang Zhang. Us-align: Universal structure alignments of proteins, nucleic acids, and macromolecular complexes. Nature Methods, 19(9):1109–1115, 2022. doi:10.1038/s41592-022-01585-1.

[45] Eric F. Pettersen, Thomas D. Goddard, Conrad C. Huang, Elaine C. Meng, Gregory S. Couch, Tristan I. Croll, John H. Morris, and Thomas E. Ferrin. Ucsf chimerax: Structure visualization for researchers, educators, and developers. Protein Science, 30(1):70–82, 2021. doi:10.1002/pro.3943.

[46] Susumu Date, Yoshiyuki Kido, Yuki Katsuura, Yuki Teramae, and Shinichiro Kigoshi. Supercomputer for quest to unsolved interdisciplinary datascience (SQUID) and its five challenges. In Sustained Simulation Performance 2021, pages 1–19. Springer International Publishing, 2023. doi:10.1007/978-3-031-18046-0_1.

